# Mining threats in high-level biodiversity conservation policies

**DOI:** 10.1101/2023.07.30.550308

**Authors:** Aurora Torres, Sophus O.S.E. zu Ermgassen, Laetitia M. Navarro, Francisco Ferri-Yanez, Fernanda Z. Teixeira, Constanze Wittkopp, Isabel M.D. Rosa, Jianguo Liu

## Abstract

Amid a global infrastructure boom, there is increasing recognition of the ecological impacts of the extraction and consumption of construction minerals, mainly as concrete. Recent research highlights the significant and expanding threat these minerals pose to global biodiversity. To what extent is this pressure acknowledged in biodiversity conservation policy? We investigate how high-level national and international biodiversity conservation policies, including the 2011-2020 and post-2020 biodiversity strategies, the national biodiversity strategies and action plans, and the assessments of the Intergovernmental Science-Policy Platform on Biodiversity and Ecosystem Services, address mining threats with a special focus on construction minerals. We find that mining appears rarely in national targets, but more frequently in national strategies with greater coverage of aggregates mining than limestone mining, yet it is dealt with superficially in most countries. We then outline an 8-point strategy to reduce the biodiversity impacts of construction minerals, which comprises actions such as targeting, reporting, and monitoring systems, the evidence-base around mining impacts on biodiversity, and the behavior of financial agents and businesses. Implementing these measures can pave the way for a more sustainable approach to construction mineral use and safeguard biodiversity.

## INTRODUCTION

Contemporary societies and economic systems are quite literally built on concrete. The key mineral components of concrete – namely, sand, gravel and limestone (hereafter *construction minerals*) – are strategic resources with environmental, social and economic values, essential for the achievement of the Sustainable Development Goals (SDGs)(Thacker et al. 2019; zu Ermgassen et al. 2019b; Torres et al. 2021; Bendixen et al. 2021). Rapid population growth, household proliferation, urbanization, and infrastructure development have accelerated their extraction in the last century (Krausmann et al. 2017) to become the most extracted solid raw materials (OECD 2018) and to account for nearly 90% of the world’s anthropogenic mass, which in 2020 outweighed all Earth’s living biomass (Elhacham et al. 2020). In an age where human activities increasingly transgress the planet’s biophysical ‘safe operating space’, the expansion of concrete infrastructure – expected to double by 2060 (OECD 2018) – comes with considerable ecological risks as a major driver of both carbon emissions and biodiversity loss (Müller et al. 2013; Torres et al. 2022 [preprint]; zu Ermgassen et al. 2022a).

The mining of construction minerals poses serious direct and indirect impacts on biodiversity through increased erosion, traffic, pollution, water stress, salinization, and land-use changes (Hughes 2017; Sonter et al. 2018; IPBES 2019; Koehnken et al. 2020). For example, limestone quarrying has been identified as the most immediate threat to karst biodiversity in Southeast Asia (Clements et al. 2006; Hughes 2017), where many species remain undescribed (Whitten 2009). The overexploitation of sand and gravel has been identified as a top priority for bending the curve of global freshwater biodiversity loss (Tickner et al. 2020). Torres et al. (2022) [Preprint] found over a thousand species in the IUCN red list reported to be threatened by mining construction minerals globally and many newly described species imminently threatened by this activity.

Despite many calls from diverse voices to pay increasing attention to this environmental pressure and scaling up solutions to the impacts of our reliance on construction minerals (Peduzzi 2014; Torres et al. 2017, 2022; CBD 2018; Hughes 2019; Tickner et al. 2020; UNEP 2022a, 2022b), it is unclear if these efforts have filtered through into conservation policy. The primary instrument for the international community’s commitment to reverse biodiversity loss over the past decade has been the United Nations’ Strategic Plan for Biodiversity 2011-2020 (Rogalla von Bieberstein et al. 2019), which was developed under the Convention on Biological Diversity (CBD), endorsed by all the biodiversity-related conventions, and adopted in 2010. Essential to the achievement of this Plan and the associated global Aichi biodiversity targets is their implementation at the national level through the formulation of national biodiversity strategies and action plans (NBSAPs) and national targets. The effective implementation of goals and targets also relies on the identification of sector-specific actions and their monitoring to promote mainstreaming, ownership, and accountability (Perino et al. 2021). In fact, the CBD has reiterated the importance of mainstreaming biodiversity within the mining and infrastructure sectors with the Sharm El-Sheikh Declaration (CBD 2018), adopted at the 14th COP meeting in 2018 (CBD/COP/DEC/14/3). However, the degree to which nations are mainstreaming biodiversity into other sectors of society varies substantially (Whitehorn et al. 2019). In December 2022, a new global framework for action on biodiversity conservation to 2030 – known as the Kunming-Montreal global biodiversity framework (GBF) – was agreed at the 15th Conference of Parties (COP15). It builds on the global assessment of the Intergovernmental Science-Policy Platform on Biodiversity and Ecosystem Services (IPBES) and its call for transformative change (IPBES 2019), as well as the Aichi targets and lessons learned from their implementation or lack thereof.

Here, we examine to what extent the mining of construction minerals is considered by high-level national and international biodiversity conservation policies. We quantify the degree to which threats from mining these minerals are addressed in 1) biodiversity goals and targets under the 2011-2020 biodiversity strategy; 2) NBSAPs and associated national targets; 3) regional and global assessments under IPBES; and 4) the newly signed Kunming-Montreal GBF. In doing so, we investigate whether and how the increased understanding on mining risks has permeated into biodiversity conservation policies. Then, we present an 8-point strategy for reducing biodiversity impacts of construction minerals mining and use.

## METHODS

To investigate the degree to which threats posed by mining construction minerals are highlighted in biodiversity conservation policies, we conducted a review of 1) the global Aichi biodiversity targets, 2) all national targets for the 2011-2020 CBD framework, 3) the latest version of all NBSAPs submitted to the CBD Secretariat, 4) the global, regional and land degradation and restoration IPBES assessments (https://ipbes.net/assessing-knowledge), and 5) the global goals and/or targets under Kunming-Montreal GBF (CBD/COP/DEC/15/4; https://www.cbd.int/doc/decisions/cop-15/cop-15-dec-04-en.pdf). Both national targets and NBSAPs in English, Spanish, French, Portuguese, and German were considered. National targets were downloaded from https://www.cbd.int/nbsap/targets/ on October-November 2022 for 176 countries, and the NBSAPs available in https://www.cbd.int/nbsap/search/ were downloaded for 181 countries (a total of 193 countries have submitted NBSAPs by November 2022). We used a text coding approach to identify the targets/documents that mentioned the topic of mining and/or specifically construction minerals mining (see Appendix S1 for the list of terms). We classified mentions into three categories: a) those referring to general threats from mining as a whole or to the need for improved planning and management of mining activities to minimize trade-offs with biodiversity conservation; b) those mentioning threats from mining construction minerals; and c) those referring to the protection of sourcing ecosystems (e.g., sandbanks, sandy beaches, limestone hills) due to their ecological significance. These categories elucidate for each target and NBSAP the relationship between biodiversity, mining, and construction sectors. We acknowledge that mentioning a threat in the national strategy does not systematically equate to implementing the proper means to address it. The recognition of these relationships in the NBSAPs is a clear indication that countries acknowledge the need to integrate biodiversity concerns into planning of the construction and mining sector although it cannot be interpreted as implementation of actions.

We examined whether mentioning construction minerals in NBSAP or national targets was associated with country-level attributes through logistic regression models with a binomial distribution and logit link function using the *glm* function from the R stats package (R Core Team 2021). We included as explanatory variables: the interaction between country size and island status based on the UN list of Small Island Developing States (https://www.un.org/ohrlls/content/list-sids); GDP per capita in the most recent year for which there was data available (all between 2017-2020) calculated using GDP and population size data from the World Bank data(World Bank 2021); average domestic extraction of construction minerals 2015-2019, calculated from the UNEP IRP Global Material Flows Database (https://www.resourcepanel.org/global-material-flows-database) using “Non-metallic minerals - construction dominant” flows; the percentage of species reported as impacted by mining construction minerals of the total number of assessed species in the red list by country from Torres et al. (2022); and the length (number of pages) of the corresponding NBSAP. The significant threshold was *P* < 0.05. We estimated maximum likelihood pseudo r^2^ using the ‘pR2′ function of the ‘pscl’ package for R (Jackman et al. 2023). We present the results of the optimal model according to the lowest Akaike’s Information Criterion, corrected for finite sample sizes (AICc) (Anderson & Burnham 2004).

Finally, we compiled policy interventions on Multi-lateral Environmental Agreements (MEAs) relevant to biodiversity conservation that include direct or indirect reference to the mining of construction minerals, from searches through policy documents, academic articles (Weyman 2016), and intergovernmental organizations reports (e.g., UNEP 2019).

## CONSTRUCTION MINERALS IN BIODIVERSITY CONSERVATION POLICY

### Aichi targets and NBSAPs

Out of the 176 parties of the CBD examined, only 15 explicitly referred to mining in their national targets (Fig. 1A; Appendix S2). Three countries, namely Fiji, Kuwait, and Nepal mentioned the extraction of sand, gravel, or limestone, while four countries (Guinea Bissau, Malaysia, Maldives, and Tajikistan) included the conservation of source ecosystems in the national targets. In contrast, the majority of NBSAPs do acknowledge the threats posed by mining to biodiversity and the environment (85.6% of the countries with available NBSAPs: 155 of 181; Fig. 1B; Appendix S3), with 45.9% specifically mentioning mining of construction minerals (83 countries of 181). Of these, sand and gravel were the most mentioned construction minerals (75 countries; 41.4% of all NBSAPs reviewed), followed by limestone (31 countries; 17.1% of all NBSAPs reviewed). Habitats from where construction minerals can be sourced were mentioned across all NBSAPs.

**Figure 1.**
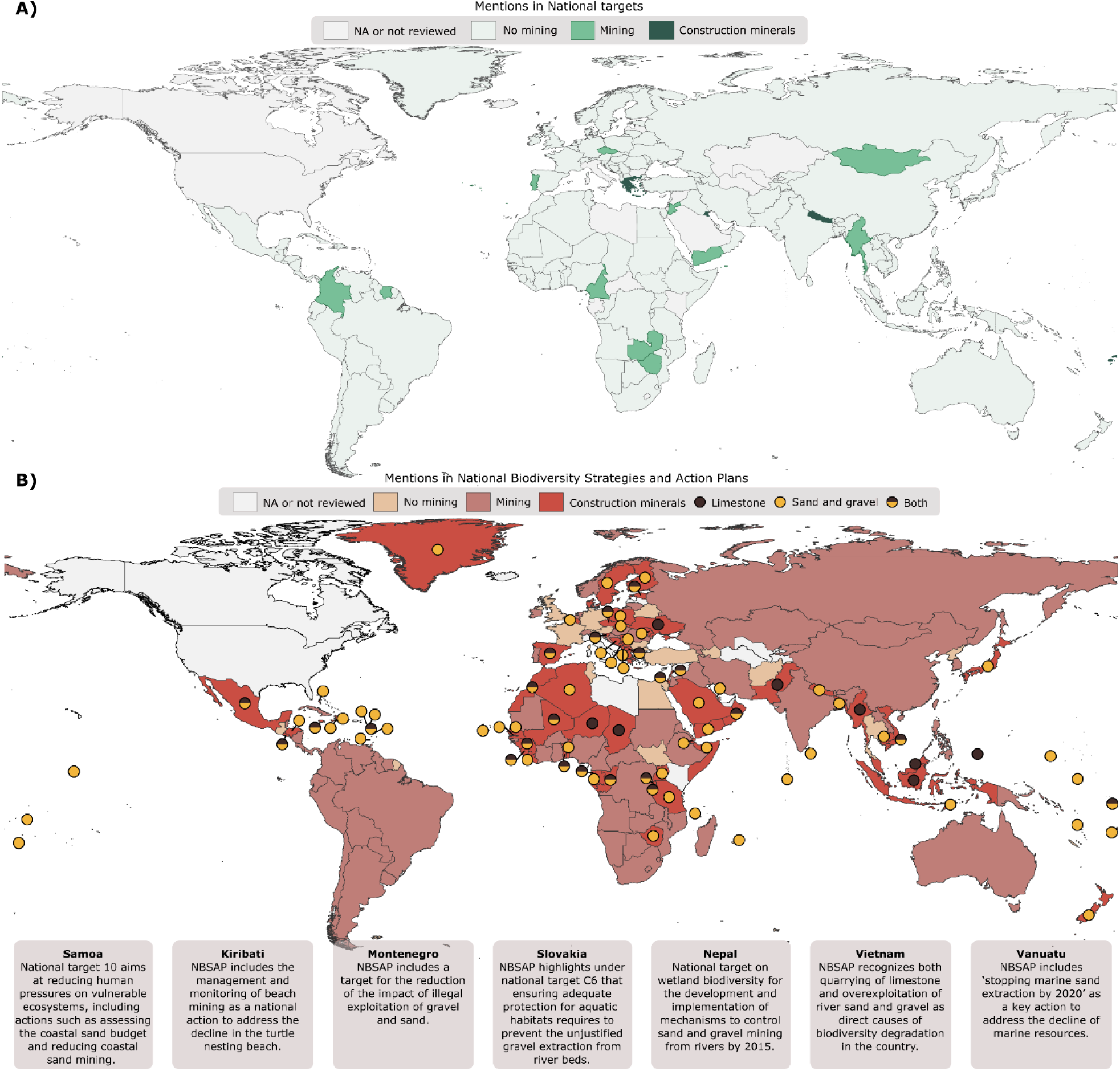
Coverage of national targets (A) and national biodiversity strategies and action plans (NBSAPs) (B) mentioning threats from and/or actions towards mining of construction minerals including a set of illustrative examples. The small round icons identify the specific type of mineral mentioned for those countries with NBSAPs that refer to construction minerals. The United States is not a party to the CBD. Full details of the targets and NBSAP mentioning construction minerals are available in Appendix S3.

The length of the NBSAPs had the most significant impact on the mentions of construction minerals, with longer assessments being more likely to address this threat (Table 1). In addition, we found that countries with a higher percentage of species impacted by mining construction minerals in the IUCN red list were more prone to adopt targets and design strategies that consider construction minerals. This was the case in countries such as Vietnam, Bangladesh, Lebanon, Malaysia, and Nepal, where the threats posed by aggregates mining and rock quarrying have been extensively documented in scientific literature and media reports (e.g., Darwish et al. 2011; Anthony et al. 2015). These findings signal the influence of the saliency of threats in the IUCN red list on the development of actions towards mining in NBSAPs and national targets.

**Table 1.**
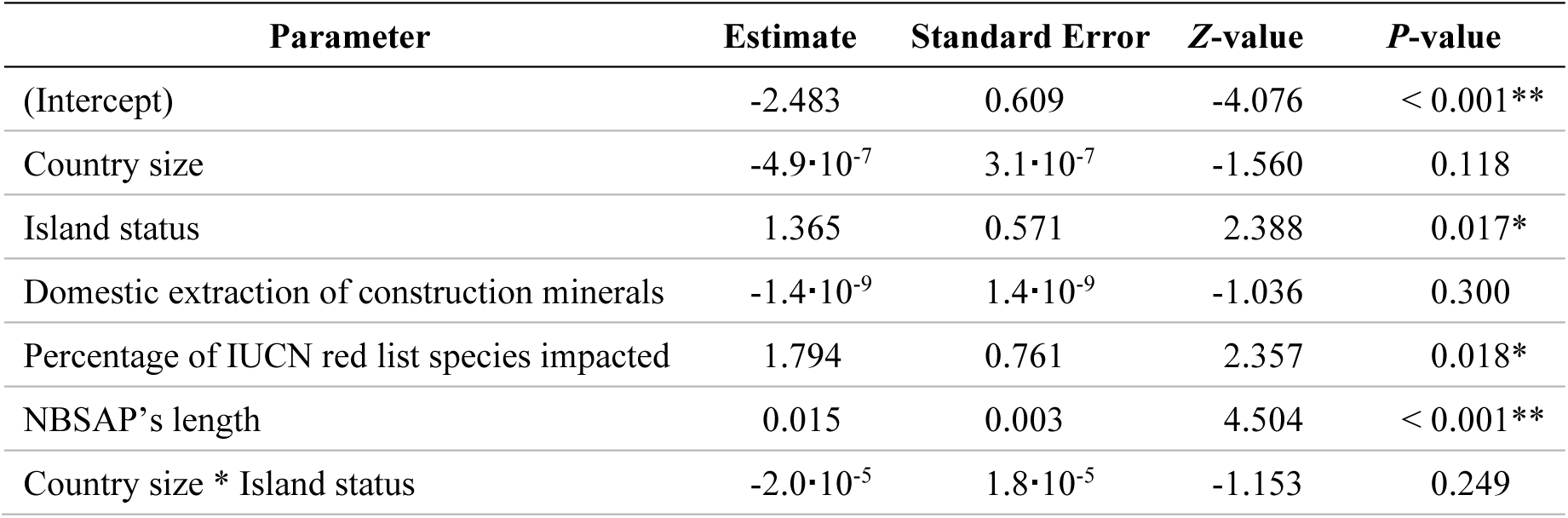
Results of the logistic regression model between including mentions of construction minerals in national targets or NBSAPs and country-level characteristics (Pseudo *R*^2^ = 0.33): country size; status of Small Island Developing States based on the UN list (https://www.un.org/ohrlls/content/list-sids); average domestic extraction of construction minerals 2015-2019, calculated from the UNEP IRP Global Material Flows Database (https://www.resourcepanel.org/global-material-flows-database); percentage of species impacted by construction minerals mining of all species assessed at the IUCN red list according to Torres et al. (2022)[Preprint]; and length of the corresponding national biodiversity strategy and action plan (NBSAP). \**P*-value < 0.05, \*\**P*-value < 0.01.

Although the interaction between country size and island status was not significant, the marginal effect of the island status was significant. This suggests that irrespective of country size, there is a higher probability of mentioning construction minerals in policy documents of small islands developing states, which might result from perceived greater risks from mining, particularly of sands (see examples in Fig. 1B). Island nations face unique environmental, social, and economic challenges. Being at the frontline of climate change impacts and natural disasters, their land, freshwater, and marine ecosystems —heavily reliant on sand resources— are critical for combatting erosion, mitigating flooding risks, and are vulnerable to biodiversity loss (UNEP 2023). Poorly planned mining can undermine the communities’ resilience and compromise mitigation and adaptation efforts, as small countries are also more susceptible to supply risks (ACP-EU 2018; Komugabe-Dixson et al. 2019).

Finally, the volume of extraction of construction minerals was not associated with mentions in national targets or NBSAPs. Countries with the highest extraction volumes, including China and India, do not directly address construction minerals in their national targets or NBSAPs, which indicates a significant reporting gap.

### IPBES Assessments

IPBES global and regional assessment reports identify mining as an industry associated with direct and indirect negative impacts on biodiversity, emissions, water quality and human health (Appendix S4). Threats from extractive activities were predominantly described in sections referring to the drivers of biodiversity change and land degradation, or to the status and trends of biodiversity and ecosystems underpinning nature’s contributions to people. Although the reports feature other minerals more prominently (e.g., gold, diamonds or coal), the global assessment on biodiversity and ecosystem services (IPBES 2019) and on land degradation and restoration (IPBES 2018) recognize 1) sand and gravel mining as an indirect driver of wetlands loss and degradation, soil erosion, and changed flood patterns, and 2) cement production as a key contributor to carbon emissions. All regional assessments considered construction minerals mining as a threat to some extent. However, it was in the Asia-Pacific assessment where the issue of construction minerals stood out prominently. The region’s rapid urbanization and industrialization and its reliance on construction minerals is described to have resulted in serious impacts on biodiversity. Those range from devastating consequences for global endemicity hotspots in karstic areas where quarrying is considered the main threat to species survival (Clements et al. 2006; Hughes 2017), to the extraction of aggregates destroying critical marine habitats such as seagrass and accelerating coastal erosion that threatens beaches and rocky shores (Thaman 2013; Peduzzi 2014; UNEP/UNCTAD 2014). In Western and Central Europe and Africa the discussion was more focused on the impacts of aggregates dredging in rivers and coastal areas, for example as an important cause of mangrove decline in West Africa. In America, the primary focus was on cement plants and rock quarries as major sources of pollution.

### Other Multi-lateral Environmental Agreements

In addition to the CBD, there are other Multi-lateral Environmental Agreements (MEAs) related to biodiversity conservation for which the extraction of construction minerals is relevant (Fig. 2; Appendix S5), including the SDGs, conventions to minimize the impact of aggregates mining on wetlands and marine areas (Radzevičius et al. 2010), and international environmental policy instruments for environmental assessment. Interestingly, we show that the saliency on the topic of the mining and use of construction minerals has increased in the international community, with three resolutions of the United Nations Environmental Assembly, and one resolution and one recommendation of the IUCN World Conservation Congress that directly address the mining of construction minerals having been adopted since 2016. It is important to note that the increased saliency of the theme in high-level biodiversity conservation policies does not necessarily translate into increased implementation efforts. Nevertheless, by design, the 2011-2020 strategic plan for biodiversity supports the mapping of targets across conventions and cooperation for their effective implementation (Rogalla von Bieberstein et al. 2019), which should be reflected in the NBSAPs and have been captured by our analysis.

**Figure 2.**
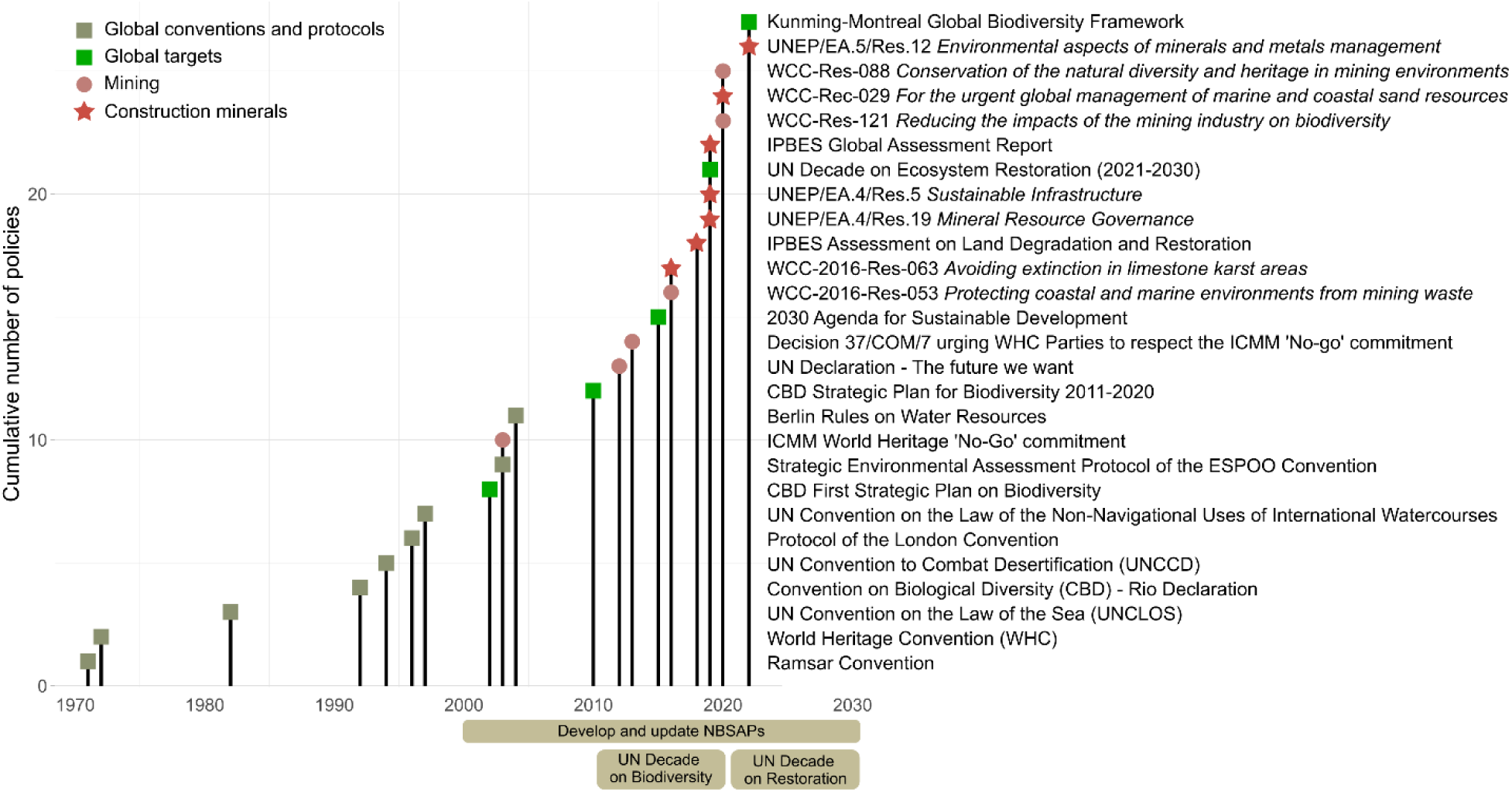
Chronology of Multi-lateral Environmental Agreements relevant for the nexus between construction minerals and biodiversity over the last 50 years. Gray squares, global conventions and/or associated protocols; green squares, global strategy and targets; dots, policy instruments that mention mining; stars, policy instruments that mention construction minerals; ESPOO, Convention on Environmental Impact Assessment in a Transboundary Context; ICMM, International Council on Mining and Metals; IPBES, Intergovernmental Science–Policy Platform on Biodiversity and Ecosystem Services; UN, United Nations; UNEP, UN Environment Program; WCC, IUCN World Conservation Congress. Since 2000 and even earlier, Parties develop and update national biodiversity strategies and action plans (NBSAPs). Full details of the listed policies and their relevance are available in Appendix S5.

### Post-2020 Global Biodiversity Framework

The Kunming-Montreal GBF aims at halting biodiversity loss, and driving its recovery, while accounting for the many benefits that humans and society can derive from healthy and sustainably used ecosystems. Three of the framework’s targets are of relevance regarding the impact of mining construction minerals. The target 12 focuses on the “green and blue spaces in cities”, “biodiversity-inclusive urban planning”, and “sustainable urbanization”, but it does not account for the off-site impacts of urban development, such as provisioning construction minerals. Target 14 on biodiversity mainstreaming into “policies, regulations, planning and development processes” would likely call for such mainstreaming within the mining sector. In fact, the draft GBF produced by OEWG4 in July of 2022 listed mining and deep-sea mining, however the text has not remain in the final version. Lastly, target 15 indicates that businesses and financial institutions must assess and report on their impacts on biodiversity and strive towards the full sustainability of their activities. The first draft of the target included “extraction practices”, but again the term was dropped. The absence of a clear reference to mining carries a risk of construction mineral mining being overlooked in future NBSAPs derived from the GBF.

## HARD PROBLEMS, CONCRETE SOLUTIONS

The Montreal-Kunming GBF is built around a theory of change that acknowledges the need for urgent policy action globally, regionally, and nationally to transform economic, social, and financial models for stabilizing biodiversity loss trends by 2030, with net improvements by 2050. However, the demand for construction minerals is projected to double by 2060 (OECD 2018), leading to mining expansion into biodiversity-rich areas (e.g., Hughes 2019). Recent assessments and resolutions stress the need for transformative changes via transitions towards sustainable pathways, including when it comes to cities and infrastructure development (Díaz et al. 2019; CBD 2020). While discussions on the “sustainable cities” transition center on green infrastructure and nature-based solutions, these documents also advocate for sustainable materials and improved spatial planning that accounts for the impact of urban communities on nearby and distant ecosystems, following the metacoupling framework (Liu 2017). Yet, the full reach of the threat posed by mining construction minerals to biodiversity remains uncertain due to knowledge gaps and deficiencies (Torres et al. 2022 [Preprint]; Cooke et al. In Press). Our analysis shows that current policies still fall short of clear statements and outcomes regarding the reporting and monitoring of mining threats, especially related to construction minerals. Moreover, international treaties require improved design for greater impact (Hoffman et al. 2022). To inform such efforts, we outline an 8-point strategy to effectively mainstream biodiversity into the extractive, infrastructure, and construction sectors (Fig. 3).

**Figure 3.**
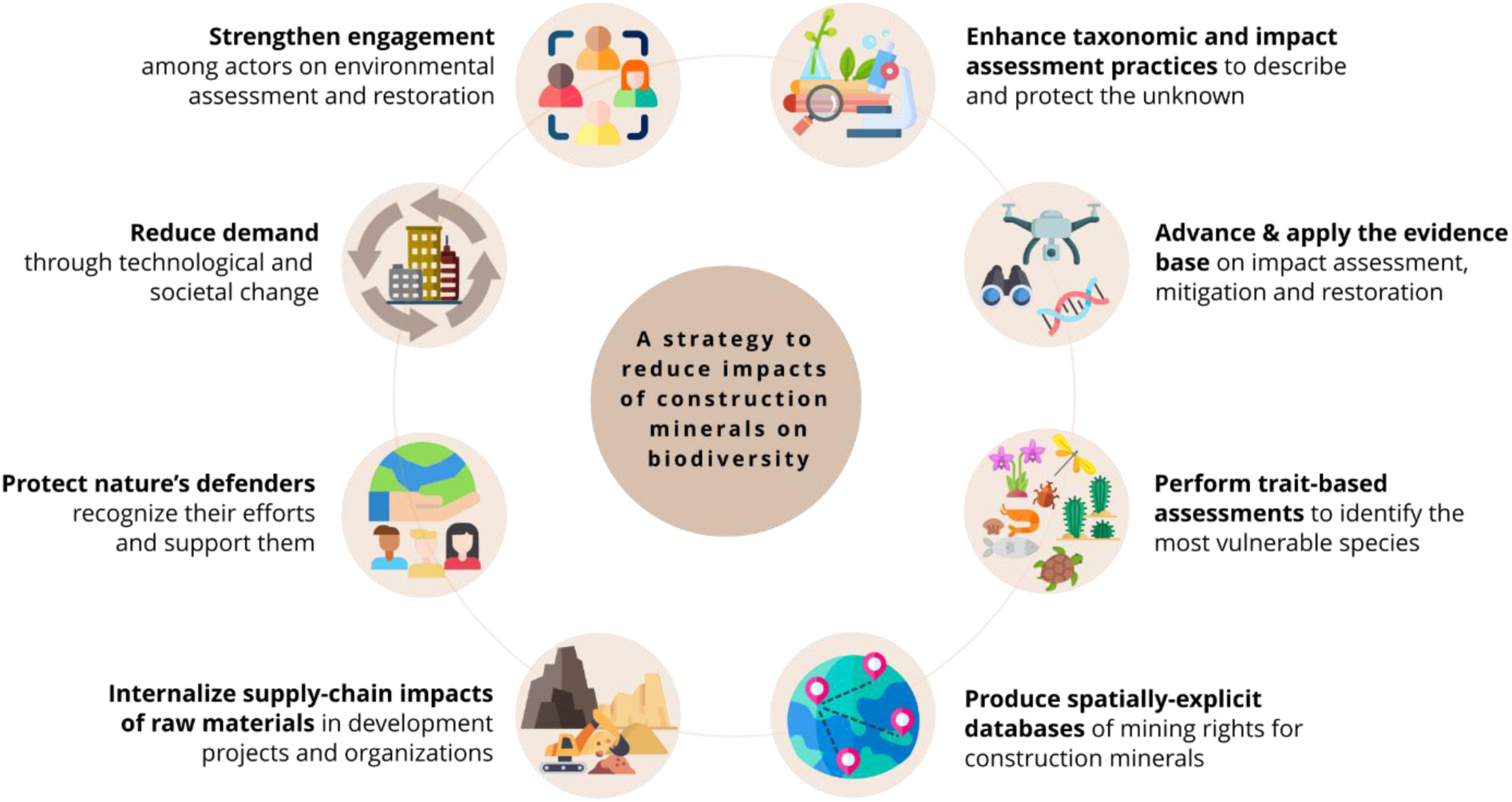
An 8-point strategy to reduce the impacts of the mining and use of construction minerals on biodiversity over time. Icons credit—https://flaticon.com.

### (1) Enhance taxonomic and impact assessment practices to describe and protect the unknown

Sound conservation decisions require knowledge of the species present. Mining construction minerals pose a threat to ecosystems that host numerous undescribed species of poorly known groups such as invertebrates, fungi, and plants (Reddy 2014; Torres et al. 2022 [Preprint]). The dire need to catalog, study, manage, and protect species and their habitats in mining frontiers clashes with a stagnation in the number of taxonomists, funding and training (Drew 2011; Sluys 2013). Bebber et al. (2014) estimated that the average lag between the collection of a plant specimen and the publication of species description was 35 years. Given the rapid development rates, even a fraction of that time would mean that many species may become extinct during the description process. Molecular approaches like DNA barcoding and metabarcoding aid in estimating biodiversity but require resources not universally available and the procedures are not explicitly designed to describe species. It is essential to ensure funding for taxonomic research and training, and foster collaboration between taxonomists and red-list assessors to provide red-list assessments as part of taxonomic descriptions (Tapley et al. 2018; Hochkirch et al. 2021). Solely prioritizing the red list, the assessment process overlooks vulnerable, unassessed, or poorly assessed species, potentially neglecting their conservation (Martín-López et al. 2011; Simmonds et al. 2020). Setting good practice in impact assessment following a risk-based approach when extractive industries enter areas with poorly documented species might address this gap: “*if a species is potentially new to science or globally threatened, has highly restricted range, and knowledge of its distribution, ecology, and of restoration needs is lacking, the precautionary principle should apply and impacts on it should be avoided. If all actors decide avoidance is unfeasible, it should not be translocated, moved or destroyed until its requirements are researched and effective techniques are available.*” (Treweek pers. Comm.)

### (2) Advance and apply the evidence base on biodiversity responses to mining and restoration

Despite decades of developments in the practice of environmental assessment and numerous guidelines (UNEP 1990; IUCN 2014), severe knowledge gaps persist regarding how to mitigate development impacts on ecosystems and restoring or offsetting biodiversity after mining (zu Ermgassen et al. 2019a; Martins et al. 2020; Christie et al. 2020; Hunter et al. 2021; Boldy et al. 2021). The current system for enhancing the evidence base is haphazard and inefficient; with the majority of post-intervention monitoring remaining unpublished, and suspected low compliance rates with mandated mitigation measures because of a lack of third-party enforcement (Tischew et al. 2010; zu Ermgassen et al. 2019a). Baseline surveys in EIA should provide transparent and evidence-based information on biodiversity impacts and mitigation recommendations (Brownlie & Treweek 2018). Maximizing the technical quality and scientific value of the follow-up monitoring of mining projects to assess the effectiveness of mitigation, restoration, and offsetting actions (e.g., integrating field surveys with environmental DNA and remote sensing) would improve the volume and efficiency of new evidence (Lindenmayer & Likens 2009; Dias et al. 2019). An ideal system for mitigating impacts and iteratively enhancing the evidence base would involve routine public reporting of monitoring outcomes according to the FAIR principles (Findable, Accessible, Interoperable and Reusable; Wilkinson et al. 2016) and open practices (e.g. published in Conservation Evidence database, mobilizing data to GBIF; King et al. 2012). This approach would contribute to the accountability of the mining sector (Perino et al. 2021). For such system to succeed, authorities must empower local institutions and organizations to access, apply, and contribute up-to-date knowledge to inform environmental assessments, decision-making, and the design of mitigation measures (UNEP 2022a). Improved and independent funding mechanisms are needed, and extractive industries should also contribute resources and data to expand the evidence base through site-based research.

### (3) Perform trait-based vulnerability assessments

In parallel to efforts to boost the reporting of mining threats on particular species, approaches based on traits (behavioral responses or life-history traits) can contribute to identify species that will be most impacted by construction minerals mining in a timely manner for conservation and management (Bland & Böhm 2016; Kopf et al. 2017; Jarić et al. 2019). This would shed light into the mechanisms that contribute to imperilment, making predictions for unassessed species, and ranking species based on their relative vulnerability. The database of species produced by Torres et al. (2022)[Preprint] can be a starting point to use as a “Robin Hood” approach (sensu Punt et al. 2011), where available assessments are used to examine species that are information-poor.

### (4) Open-access maps of mining rights for construction minerals

Mining rights for aggregates or limestone are barely represented in global mining databases (SNL Metals and Mining or S&P Global Market Intelligence databases) or land-cover datasets, and insufficiently covered by many national datasets. The lack of comprehensive mapping is concerning as it might easily downsize the environmental and social risks posed by mining. Producing publicly available spatially-explicit databases of mining rights for construction minerals would be a major step to understand the extent and distribution of biodiversity threats and for identifying opportunities for mine restoration. Along with that there is a need for greater appreciation and improved characterization of the diversity of mining contexts (Sonter et al. 2022). Mining hotspots, industry structure, regulatory frameworks, and supply chains differ considerably among minerals (Franks 2020). However, many studies refer to mining as land-use without reporting the extracted material (Boldy et al. 2021), an issue that also affect about 20% of the red list assessments referring to mining threats (Torres et al. 2022 [Preprint]). To maximize the usage of assessments, the characterization of mining threats should include information on the minerals type, geographic location, mining methods, mining intensity, and the impact mechanisms.

### (5) Account for supply-chain impacts of raw materials when financing development projects and assessing organizational biodiversity footprints

Including the impacts of mining construction minerals and their supply chains within the scope of multilateral and private finance environmental safeguard policies would internalize the ecological costs of extraction. As it stands, major multilateral development banks’ safeguard policies hold their clients responsible for some supply-chain impacts of the projects they help finance, but often non-living raw materials are excluded (Table 2). A simple wording change, adopting the World Bank safeguards’ definition of raw materials (which explicitly includes sand and other construction minerals), could be a valuable leverage point, laying the groundwork for internalizing the supply-chain impacts of construction minerals consumption into tens of billions of dollars’ worth of project financing each year. Likewise, financial institutions need to assess their exposure to environmental risks associated with investments reliant on dredging marine aggregates (e.g., land reclamation projects) as highlighted by UNEP Finance Initiative (UNEP 2022b). As organizations and international institutions also strive to deliver ‘nature-positive’ outcomes, there is a growing focus on addressing supply-chain impacts through organizational sustainability strategies (zu Ermgassen et al. 2022b), since the majority of impacts for businesses are embedded in those chains, not their direct operations (e.g., Bull et al. 2022). The little work that has been done reveals substantial impacts of construction mineral supply chains. In an analysis of the University of Oxford’s biodiversity footprint, the biodiversity impacts and emissions embedded in construction supply chains were one of the largest categories of the organization’s impacts, with construction and cement use ranking as major drivers of water consumption, acidification, and eutrophication (Bull et al. 2022). However, methodological gaps remain, with footprinting largely relying on impact estimates averaged across a bundle of related economic activities (e.g., those in databases like Exiobase) and lacking spatial considerations.

**Table 2.**
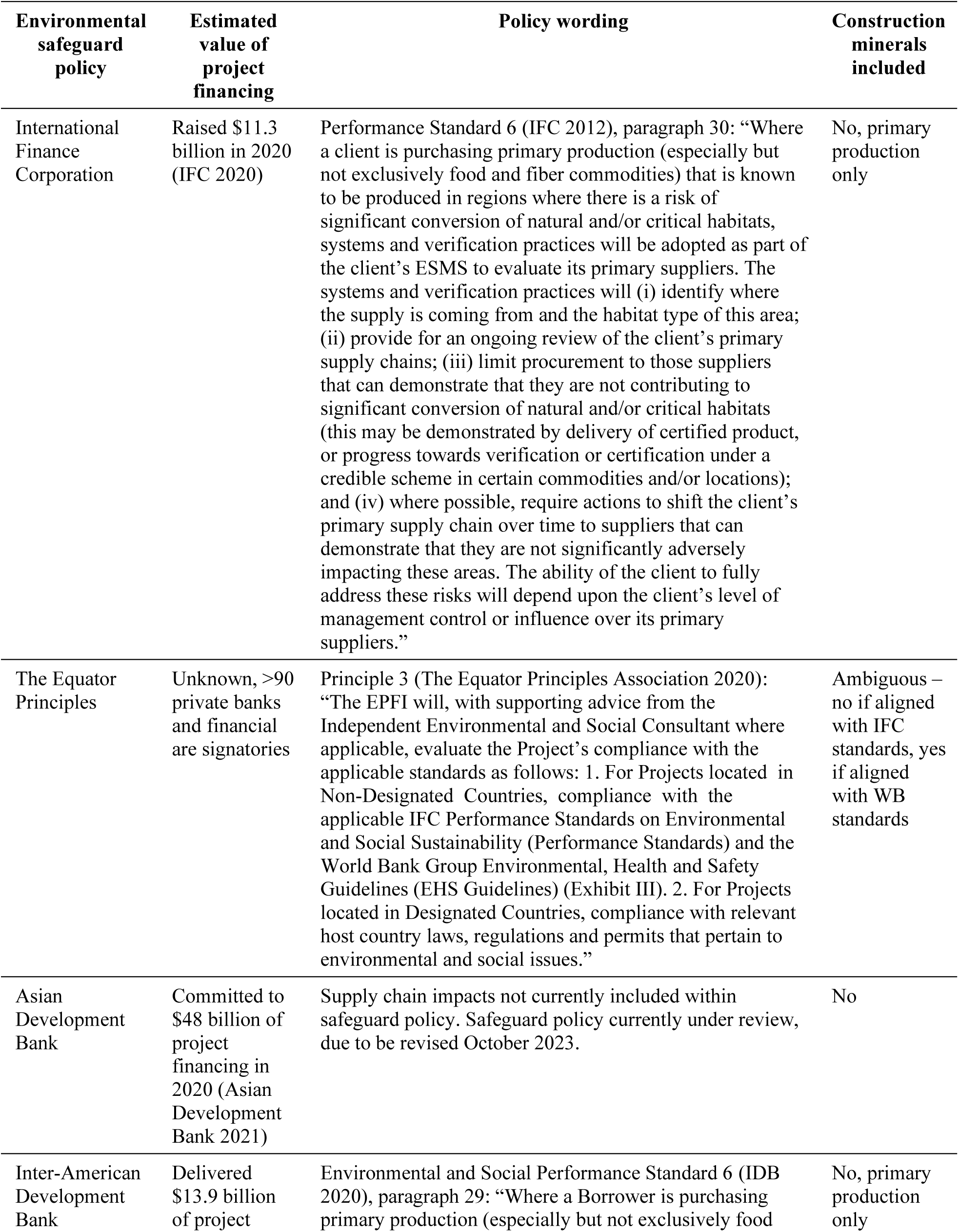
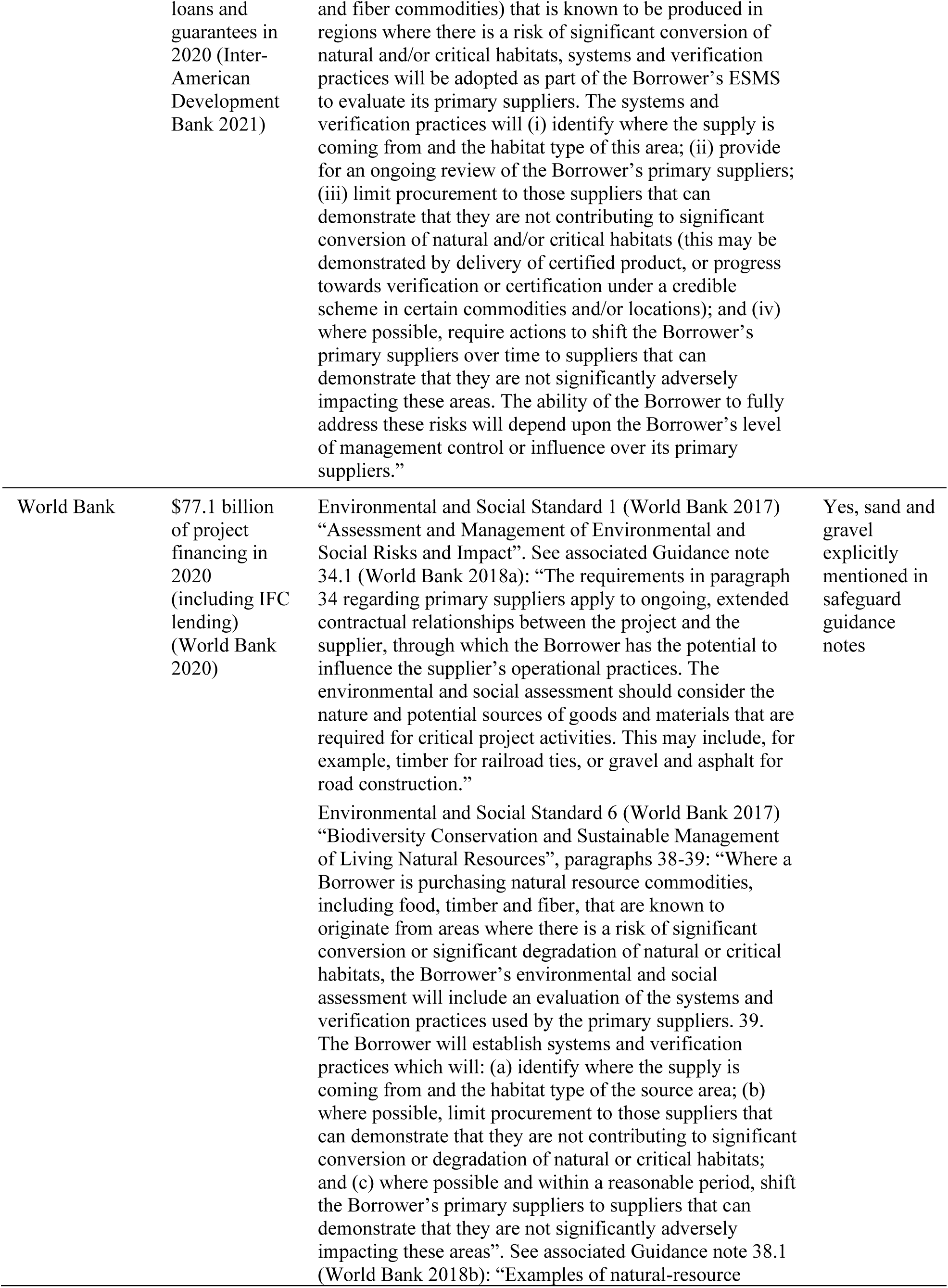
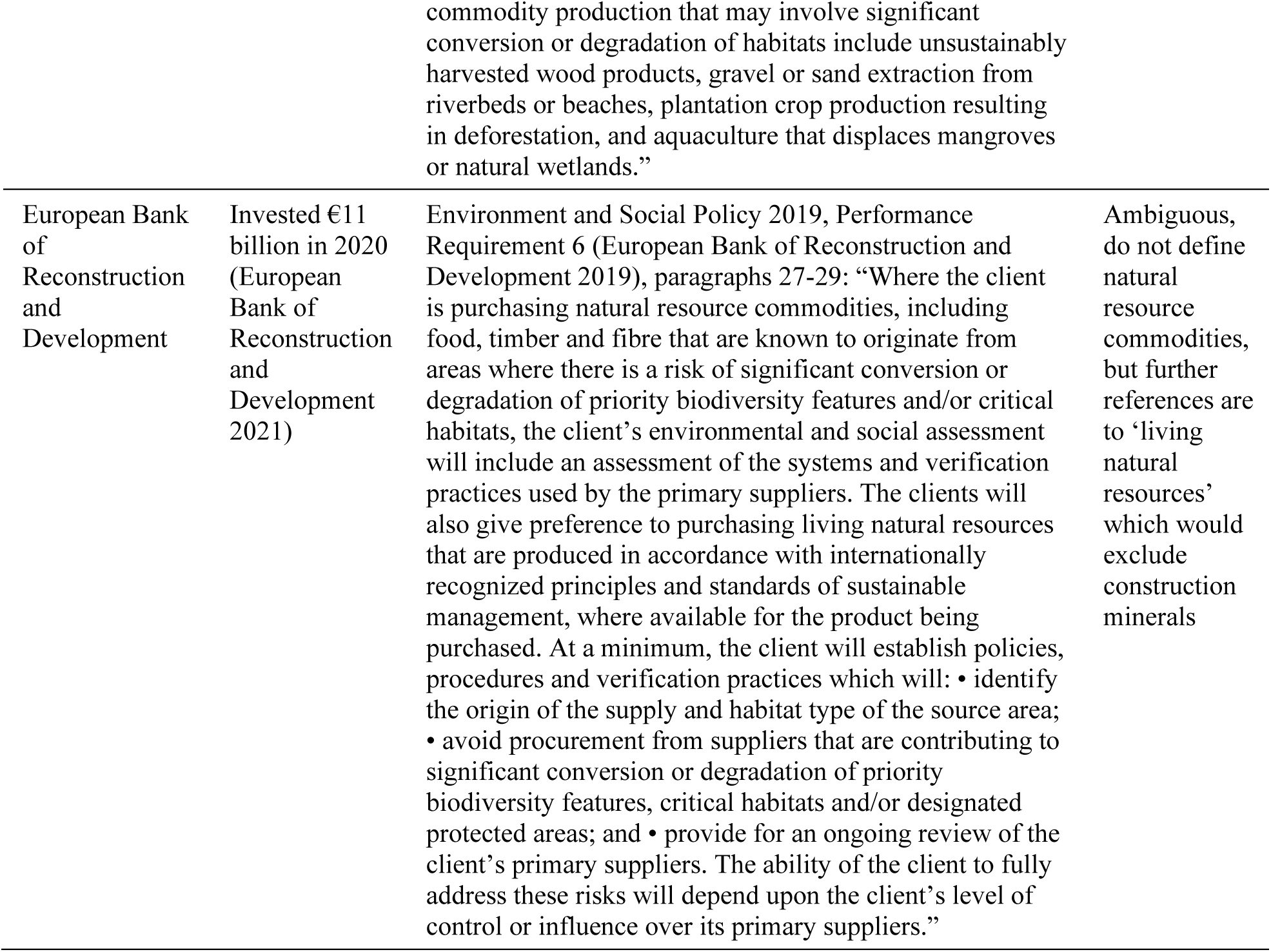
Coverage of construction minerals in major multilateral development banks’ safeguard policies.

### (6) Protect nature’s defenders

Target 22 of Kunming-Montreal GBF recognizes the rights of Indigenous Peoples and local communities and emphasizes the need to ensure the participation in decision-making, and the full protection of environmental human right defenders. Local civil society organizations and community groups, at the forefront of conservation efforts, often face threats and violence while protecting nature from unsustainable mining (Glazebrook & Opoku 2018; Bebbington et al. 2018). The murder of land and environmental defenders is a widespread and growing phenomenon (Zeng et al. 2021), particularly affecting the mining sector, which has reported the highest number of murders (Global Witness 2020). Instances of illicit activities and violence related to sand and limestone mining are common in various countries, including Mexico, Turkey, Kenya, India, China, Indonesia, and Cambodia (Constable 2017; REFORMA 2019; SANDRP 2019; Bisht 2021). This situation undermines meaningful engagement among stakeholders and inhibits the establishment of a cohesive community of practice. Without ensuring the safety of nature defenders, it becomes nearly impossible to gather accurate information on the impacts and risks to biodiversity from mining construction minerals. Urgent government protection, local support, international recognition, and the mobilization of human-rights mechanisms are needed to address these issues and underlying factors (Glazebrook & Opoku 2018; Bille Larsen et al. 2021).

### (7) All the previous steps are likely insufficient on their own without addressing the rapid growth in demand for construction minerals

Global material stocks are projected to increase by 66% from 2015-2035, despite scientists warning that the global economy is already consuming materials in excess of that required to remain within Earth’s ‘safe-operating space’ (Bringezu 2015; Wiedenhofer et al. 2021). Haberl et al. (2019) show that there is a non-linear relationship between national concrete stocks and material improvements in people’s wellbeing, with a satiation point around 50 t concrete/capita, suggesting that increasing concrete stocks in infrastructure-rich nations may be unnecessary for meeting people’s fundamental needs. There is increasing recognition that society experiences high-carbon lock-in effects at least partly because of an overriding political economy that favors high-resource consumption pathways to meet societal demands (these political-economic high-carbon lock-ins have been reviewed for the automobile and housing sectors: Mattioli et al. 2020; zu Ermgassen et al. 2022a). Addressing these and reducing materials demand is an essential component of achieving sustainable levels of construction mineral mining and consumption (Creutzig et al. 2018; Bisht 2022). This requires rapid rates of innovation-driven de-materialization coupled with substantial changes in economic systems, such as making more efficient use of existing infrastructure instead of satisfying further demand solely through infrastructure expansion (IRP 2019; Zhong et al. 2022).

### (8) Strengthen engagement among sectors to create a community of practice

The International principles and standards for the ecological restoration and recovery of mine sites highlight that the best environmental outcomes are achieved when engagement and trust are established among extractive companies, government agencies, scientists, and local communities (Young et al. 2022). These principles advocate for a *Trust Model*, promoting genuine, open, and transparent interactions amongst sectors, which are essential for obtaining and maintaining the social license to operate, with science providing independent oversight for quality assurance. Recent industry initiatives on the construction minerals sector align with the principles of the UN Decade for Restoration, seeking to inspire positive change (e.g., CEMBUREAU 2022). Various cases show that collaborative research and engagement are crucial for establishing meaningful conservation and restoration targets, defining priorities to allocate resources, and pinpointing implementation obstacles and knowledge gaps (Rokich 2016; BirdLife Europe and Central Asia & HeidelbergCement 2017; Salgueiro et al. 2020). Long-term relationships between mining and research projects can strengthen the knowledge base by using powerful study designs (e.g., before-after-control-impact designs or randomized experiments), thereby increasing the inferential strength of assessments and informing effective strategies along the mitigation hierarchy. Such efforts will help identify what works and under which conditions, and how efforts can be scaled up, shared, and studied. However, research institutions must be careful not to legitimize malpractice – research funds are no substitute for impact avoidance when mining impacts on threatened or poorly known biodiversity.

## CONCLUSIONS

Following the adoption of the Montreal-Kunming GBF, countries will develop or revise their national biodiversity strategies. We encourage the adoption of the proposed strategy for reducing the biodiversity impacts of mining construction minerals over time. Initiatives aimed at addressing data and knowledge gaps will help iteratively improving the scientific knowledge that underpins policies governing mineral resources through international treaties, national and subnational policies and strategies, and platforms. A wide network of local-to-global partners covering environmental conditions and socioecological contexts is essential. Specific actions must be taken to set up targets to monitor mining threats, enhancing transparency, and informing policies across sectors such as nature conservation and restoration (e.g., SDG 14 and 15, UN Decade on Ecosystem Restoration), and urban sustainability (SDG 11). Implementing the recommended measures will contribute to securing the social license to operate, empowering industry, authorities, and civil society to cultivate stronger relationships that drive systemic improvements throughout industry and hold key stakeholders accountable. These actions must be part of a wider transformative change to transition to less resource-intensive economies for addressing society’s infrastructure needs.

## SUPPORTING INFORMATION

Appendix S1-S5

## ACKNOWLEDGEMENTS

This article has benefited from constructive comments and helpful suggestions from J. Treweek, J. Simmonds, and E.F. Lambin. A.T. received funding from the European Union’s Horizon 2020 research and innovation programme under the Marie Sklodowska-Curie grant agreement No 846474 and the Generalitat Valenciana (CIDEIG/2022/44). L.M.N and F.F. are funded by the Spanish Ministry of Science and Innovation project SUMHAL (LIFEWATCH-2019-09-CSIC-13). F.Z.T. is funded by Coordenação de Aperfeiçoamento de Pessoal de Nível Superior (PNPD/CAPES, Finance Code 001). J.L. is supported by the National Science Foundation and Michigan AgBioResearch. This article contributes to the objectives of the Global Land Programme (https://glp.earth).

