## Appendix S1 for "Mining threats in high-level biodiversity conservation policies"

**Appendix S1**. Search terms considered for the identification of relevant national targets and biodiversity strategies and action plans in multiple languages.

| English | Spanish | French | Portuguese | German |
|---|---|---|---|---|
| mineral | mineral | minerai / minéraux / minéral | mineral / minerais | mineral |
| mining | minería | exploitation minière / mine | mineração | abbau |
| quarr | minería | carrière | pedreira | abbauen |
| dredg | draga | drag (dragage) | dragagem | ausbaggern |
| aggregate | áridos / agregados | | | aggregat |
| sand | arena | sable | areia | sand |
| gravel | grava | gravier | gravilha / cascalho | kies |
| stone | | moellon | | stein |
| limestone | caliza | calcaire / moellon | calcário | kalkstein |
| cement | cemento | ciment | cimento | zement |
| concrete | | | | beton |
