## Appendix S2 for "Mining threats in high-level biodiversity conservation policies"

**Appendix S2**. Detailed search results on Aichi national 2011-2020 targets for countries that mentioned at least one relevant search term of mining or construction minerals (see Appendix S1 for a full list of search terms in multiple languages). In bold red the search term identified in the target's quote.

| **Considers explicitly** | | | | | |
|---|---|---|---|---|---|
| **Mining and/or dredging** | **Extraction of construction minerals*** | **Source ecosystems** | **Country** | **National target name** | **Target text** |
| x | | | Cameroon | E-Target 4 | Develop and/or intensify integrated action frameworks on all activities (**mining**, industrial logging, smallholder agriculture, and illegal logging) that impact on forest biodiversity conservation, Protected Areas management in a manner that enhances local governance. |
| x | | | Colombia | METAS 2020, III.10 | Se habrá establecido un sistema de fiscalización y de rendición de cuentas de los efectos ambientales de las actividades productivas relacionadas con **minería**, hidrocarburos, infraestructura, ganadería y agricultura. |
| | | | | METAS 2030, V.1 | A 2030 (…) Se habrán controlado los principales motores de pérdida y degradación de bosques en el país: ampliación de la frontera agrícola; colonización asociada a pastos para la ganadería, **minería**, incendios forestales, cultivos de uso ilícito; infraestructura (centros urbanos y construcción de vías) y extracción de madera. |
| | | | | METAS 2025, III.10 | III.8 El 100% de los proyectos de concesión de infraestructura de 4G, programas de desarrollo **minero** y expansión energética, vivienda y ciudades amables, agricultura y desarrollo rural contarán con evaluaciones ambientales estratégicas. |
| | | | | METAS 2025, III.10 | III.9 Los sectores económicos agropecuario, **minero** energético e infraestructura contarán con indicadores de sostenibilidad y con mecanismos de seguimiento y verificación del cumplimiento. |
| x | | | Czech Republic | National Target 3.4 | Soil and **mineral** resources |
| x | x | x | Fiji | Fiji Target | The NBSAP strategies and action plans are fully incorporated into 5 & 20 Year National Development Plan, the Green Growth |

| | | | | | |
|---|---|---|---|---|---|
| | | | | | Framework and other sectoral plans (e.g. Renewable Energy, Agriculture, Forestry, **Mining**, Tourism, etc.) |
| | | | | Objective SUD1 | Integrating biodiversity conservation and sustainable use and management into the production and manufacturing downstream processing and value adding for agriculture, fisheries, forestry, tourism, **mining**, other land-uses and transport industries. |
| | | | | Strategic Area SUD3 | Reducing major threats to forest and freshwater ecosystems from unsustainable logging, agriculture, fisheries, **mining** and human settlements. |
| | | | | Strategic Area SUD4 | Reducing major threats to inland waters (watershed, streams, rivers and lakes) such as **dredging**, floods, **gravel** extraction, **mining**, agriculture, deforestation, tourism, sugar, manufacturing, waste management. |
| | | | | Strategic Area SUD5 | Reduce major threats to Fiji's coastal ecosystems such as reclamation, unsustainable tourism development, river **dredging** and pollution. |
| x | | | Greece | Specific Target 5.7 | Ensure the compatibility of **mining** activities with biodiversity conservation. |
| | | x | Guinea-Bissau | National Goal 10 | By the year 2020, to identify the multiple anthropogenetic pressures on the mangroves, mud and **sand** banks and, moreover, marine and coastal ecosystems affected by the climate change or oceanic acidification and to establish strategies and programs so that their integrity and operation are maintained. |
| x | | | Jordan | National Target 12 | By 2016, renewable energy and **mining** strategies are reviewed and biodiversity safeguards adopted and enforced. |
| | x | | Kuwait | General measures medium term action plan | Use of biological resources (actions include, among others, the review of current agricultural support programs, with a view to identifying government subsidies that have adverse effects on biodiversity, the re-establishment of the land tenure allocation system and the establishment of rights of stakeholders to use the land at fair prices which will encourage water protection; preparation and implementation of a national strategy for the protection of health; introduction of a system to address problems linked to overgrazing, **sand** movement and stone clearance) |

| | | | | | |
|---|---|---|---|---|---|
| | | x | Malaysia | Target 7 | By 2025, vulnerable ecosystems and habitats, particularly **limestone** hills, wetlands, coral reefs and seagrass beds, are adequately protected and restored. |
| | | X | Maldives | Target 18 | By 2025, at least 10% of coral reef area, 20% of wetlands and mangroves and at least one **sand** bank and one uninhabited island from each atoll are under some form of protection and management. |
| x | | | Mongolia | Goal 12 | Create a legal environment where subsidies or financial assistance are prohibited for use in agriculture, **mineral resource extraction, infrastructure**, energy, light industry, food manufacturing, and service industry projects and actions deemed to be harmful to or potentially harmful to biological diversity in accordance with environmental strategy evaluations. |
| x | | | Myanmar | Target 4.1 | By 2020, SEA conducted and guidelines prepared for **mining** and energy sectors. |
| x | x | | Nepal | Target on wetland biodiversity | An effective mechanism to control **mining** of **gravel** and **sand** from rivers and streams developed and implemented by 2015. |
| x | | | Portugal | Objectivo 3.7 | Assegurar a conservação da biodiversidade e da geodiversidade nas atividades de prospeção, pesquisa e exploração de recursos **minerais**. |
| x | | | Suriname | Sub-objective 1.4 | Responsible **mining** with minimisation of damage to the environment and biodiversity and environmental restoration. |
| | | x | Tajikistan | Target 11 | By 2020, at the latest: To improve and strengthen preservation and rational use of biodiversity in order to ensure optimal provision of ecosystems services, particularly, high-mountain cryophyte, low-mountain **sand**-desert ecosystems, xerophyte light forest ecosystems, savannah ecosystems. Therewith, mesophile broad-leaved walnut ecosystems have top priority. |
| x | | | Yemen | Target 11 | By 2025, several business communities and public sectors, including ecotourism, **mining**, energy, industry and land use planning are benefiting from ecosystem services and have incorporated sustainability & biodiversity concerns into their |

| | | | |
|---|---|---|---|
| | | | national and local development plans and programmes, keeping the impacts of use of natural resources well within safe ecological limits (Aichi target 4). |
| x | Zambia | Target 8 | By 2020, pollution, including excess nutrients from industry (**mining**, agriculture, etc.), has been brought to levels that are not detrimental to ecosystem function and biodiversity. |
| x | Zimbabwe | Target 2 | By 2020, biodiversity has been mainstreamed into all seven sectors (**mining**, agriculture, health, manufacturing, transport, energy and tourism) and incorporated into national accounting and reporting systems. |
