## Appendix S3 for "Mining threats in high-level biodiversity conservation policies"

**Appendix S3**. Detailed search results on mentions of relevant search terms of mining or construction minerals (see Appendix S1 for a full list of search terms in multiple languages) in national biodiversity strategies and action plans (NBSAPs).

| **Considers explicitly** | | | | | |
|---|---|---|---|---|---|
| **Mining and/or dredging** | **Construction minerals mining** | **Construction mineral mentioned** | **Country** | **Document name** | **Details and example mention (if relevant)** |
| No | No | | Afghanistan | NATIONAL BIODIVERSITY STRATEGY & ACTION PLAN | |
| Yes | Yes | Sand and/or gravel | Albania | Document of Strategic Policies for the Protection of Biodiversity in Albania | The removal of sand and gravel for the construction industry is highlighted as a cause of coastal erosion (p. 78). |
| Yes | Yes | Sand and/or gravel | Algeria | Stratégie et plan d'action nationaux pour la biodiversité 2016-2030. | Sand extraction as a cause of loss and degradation of coastal habitats (p. 39). |
| No | No | | Andorra | ESTRATEGIA NACIONAL DE LA BIODIVERSIDAD DE ANDORRA (ENBA) | |
| Yes | No | | Angola | NATIONAL BIODIVERSITY STRATEGY AND ACTION PLAN 2019-2025 | |
| Yes | Yes | Sand and/or gravel | Antigua and Barbuda | Antigua & Barbuda National Strategic Biodiversity Action Plan 2014-2020 | Mining and dredging of sand as a cause of habitat loss and impacts on sea turtle populations (p. 12-13). |

| | | | | | |
|---|---|---|---|---|---|
| Yes | No | | Argentina | ESTRATEGIA NACIONAL DE BIODIVERSIDAD 2016-2020 | Mining has a significant impact on the biodiversity of mountanous regions (p. 9). |
| Yes | No | | Armenia | Strategy of the Republic of Armenia on Conservation, Protection, Reproduction and Use of Biological Diversity | Open mining is mentioned as a cause of habitat loss (p. 11). The Strategic direction 3 "Reduction of direct pressures on biodiversity and promotion of sustainable use" includes a key action (3.1) to "Assess the impact of small hydropower plants and mining industry on biodiversity and ESs, develop and implement an action plan on impact elimination/mitigation" (p. 28). |
| Yes | No | | Australia | Australia's Biodiversity Conservation Strategy 2010 –2030 | Mainstreaming biodiversity in the mining sector is a priority for action. However, the newer strategy for 2019-2030 does not mention mining neither other relevant terms and it is very brief (38 pages): Australia's Strategy for Nature 2019-2030 - Australia's national biodiversity strategy and action plan https://www.cbd.int/doc/world/au/au-nbsap-v3-en.pdf |
| No | No | | Austria | BIODIVERSITY STRATEGY AUSTRIA 2020+ | |
| No | No | | Azerbaijan | National Strategy of the Republic of Azerbaijan on Conservation and Sustainable Use of Biodiversity for 2017-2020 | Neither the current nor the previous NBSAPs include references to mining. |
| Yes | Yes | Sand and/or gravel | Bahamas | National Biodiversity Strategy and Action Plan | Rock and sand mining and dredging are "Priority Issues Relevant to Resource Management" (p. 72) as they have the potential for serious inter-related conflicts over the allocation and use of natural resources. "Rock and Sand Mining: There will be increasing pressure, in the years ahead, from the Eastern Seaboard States of the United States, for Bahamian sand. Considering this, the Government should seek to determine "sand cells" where commercial extraction is economically viable and environmentally sound." "Dredging: |

| | | | | | |
|---|---|---|---|---|---|
| | | | | | Government is promoting investment in marinas and cruising facilities for residents and tourists. The capacity to evaluate and monitor potential impacts of such activities is required, in order to mitigate serious environmental and health problems resulting from habitat alteration and pollution." In this case, rock mining refers to corals. |
| Yes | Yes | Sand and/or gravel | Bahrain | The National Biodiversity Strategy and Action Plan | Amongst the regulatory measures govering biodiversity, sand excavation is mentioned with the objetctive "Ban extraction and export" (p. 37). Dredging is highlighted as a cause of loss of seagrass beds (p. 41) and there is a national priority action to "Cease dredging and landfill in significant areas" in marine and coastal systems (p. 44). |
| Yes | Yes | Sand and/or gravel | Bangladesh | NATIONAL BIODIVERSITY STRATEGY AND ACTION PLAN OF BANGLADESH 2016-2021 | Unplanned sand collection from rivers is listed as a threat to biodiversity in Bangladesh (p. 21). |
| No | No | | Barbados | REPORT: NATIONAL BIODIVERSITY STRATEGY AND ACTION PLAN 2020 | NBSAP highlights that "there is a noticeable trend in the involvement of the private sector in conservation and biodiversity maintenance e.g. the conversion of quarry mines to ecologically balanced spaces". |
| No | No | | Belarus | National Action Plan for the Conservation and Sustainable Use of Biodiversity for 2016-2020 | |
| Yes | Yes | Sand and/or gravel | Belgium | Biodiversité 2020, Actualisation de la stratégie nationale de la Belgique | Impacts of the extraction of sand and gravel in river and coastal systems are highlighted (p. 24). |
| Yes | No | | Belize | NATIONAL BIODIVERSITY STRATEGY AND | Dredging of seagrass is identified as a cause of land-use change. Climate change could increase the dredging activity "for landfill as rising sea inundates cayes and coastline" (p. |

| | | | | | |
|---|---|---|---|---|---|
| | | | | ACTION PLAN (2016-2020) | 50). Although it mentions dredging it does explictly mentions marine aggregates extraction nor connects dredging to construction projects. |
| Yes | Yes | Sand and/or gravel | Benin | Stratégie et Plan d'Action pour la Biodiversité 2011-2020 | Mining of sand and gravel is recognized as a threat to the biodiversity of river and coastal systems (p. 36). |
| Yes | No | | Bhutan | NATIONAL STRATEGIES AND ACTION PLAN | Pressures from mining on biodiversity through habitat loss and degradation are highlighted (p. 53). |
| Yes | No | | Bolivia | National Biodiversity Strategy and Action Plan (v.2) | NBSAP includes a Target for 2025 for the adequate treatment of biodiversity in mines ("Desarrollado el protocolo de actuación para el adecuado tratamiento de la biodiversidad en minería") (p. 93). |
| Yes | Yes | Sand and/or gravel | Bosnia and Herzegovina | Strategy and Action Plan for Protection of Biological Diversity in Bosnia and Herzegovina (2015-2020) | Within Aichi Target 5 (Natural habitats) the NBSAP highlights that "excessive and uncontrolled exploitation of sand, gravel and other river materials leads to changes in the regime of surface and ground water, resulting in the destruction of habitats of plant and animal species in lower river courses, primarily of Rivers Bosna and Drina, and in some parts of the Rivers Sava, Vrbas and other rivers" (p. 44). |
| Yes | No | | Botswana | National Biodiversity Strategy and Action Plan | Mention of soda ash mining. One of the pressure indicators is "Proportion of each protected area boundary that has become fixed and impermeable due to settlement, lands, fencing or other land uses (e.g. mining)" (p. 121). |
| Yes | No | | Brazil | Estratégia e Plano de Ação Nacionais para a Biodiversidade | Within Target 6 the Action 13 indicates that the impacts of mining on fish fauna through the dredging of rivers must be assessed (p. 162). |
| No | No | | Brunei Darussalam | National Biological Resources (Biodiversity) Policy and Strategic Plan of Action | |

| | | | | | |
|---|---|---|---|---|---|
| No | No | | Bulgaria | National Biodiversity Conservation Plan for 2005-2010 | |
| Yes | No | | Burkina Faso | PLAN D’ACTION NATIONAL 2011-2015 DU BURKINA FASO POUR LA MISE EN ŒUVRE DE LA CONVENTION SUR LA DIVERSITE BIOLOGIQUE | Mining is mentioned as part of the 2009 national development strategy: "l’exploitation rationnelle des ressources minières" (p. 10). |
| Yes | Yes | both | Burundi | Stratégie Nationale et Plan d’Action sur la Biodiversité 2013-2020 | Mining of sand and limestone are highlighted as causes of loss and degradation of aquatic biodiversity (p. 54). |
| Yes | Yes | Sand and/or gravel | Cabo Verde | National Biodiversity Strategy and Action Plan | The extraction of "inerts" is mentioned as a cause of destruction and/or degradation of terrestrial and marine habitats and specifically it highlights the challenge of illegal exploitation of sand resources: "Although Decree-Law No. 2/2002 prohibits "the extraction and exploitation of sand dunes, beaches and inland waters, coastlines and the territorial sea", there has been a progressive increase in the consumption of sand after the approval of the creation Decree-Law. This shows the inefficiency of the Decree-Law in addressing the problem of illegal exploitation of inert (Lopes, 2010)" (p. 42-43). |
| Yes | Yes | Sand and/or gravel | Cambodia | NATIONAL BIODIVERSITY STRATEGY AND ACTION PLAN | The Strategic Objective 2 (Identify ways and means to address environmental security through preventive, proactive and corrective measures) includes a Key Action on "Prevent the damage that may occur as a result of flooding, drought, watershed degradation, erosion and sedimentation to protect fish resources and other aquatic biodiversity. These measures can include the construction or rehabilitation of bank protection works, dikes, and the provision of water storage facilities; the prohibition of sand mining on the bed and |

| | | | | | |
|---|---|---|---|---|---|
| | | | | | banks of water bodies or of the obstruction of flow/drainage, with the participation of communities (p. 46). |
| Yes | No | | Cameroon | NATIONAL BIODIVERSITY STRATEGY AND ACTION PLAN VERSION II (NBSAP II) | The Ecosystem specific Target 4 explicitly mentions mining: "Develop and/or intensify integrated action frameworks on all activities (mining, industrial logging, smallholder agriculture, and illegal logging) that impact on forest biodiversity conservation, Protected Areas management in a manner that enhances local governance" (p. 91). |
| Yes | No | | Central African Republic | STRATÉGIE NATIONALE POUR LA STRATÉGIE NATIONALE POUR LA CONSERVATION DE LA DIVERSITÉ BIOLOGIQUE EN RÉPUBLIQUE CENTRAFRICAINE | |
| Yes | Yes | Limestone | Chad | STRATEGIE NATIONALE ET PLAN D'ACTIONS SUR LA DIVERSITE BIOLOGIQUE 2ème édition | The strategy highlights that Chad has significant minerals resources (including cement) whose exploitation has negative impacts on natural resources (e.g., wildlife, flora, water, soil) (p. 50). |
| Yes | No | | Chile | ESTRATEGIA NACIONAL DE BIODIVERSIDAD 2017-2030 | Within the Strategic Objective "Introduce biodiversity objectives in policies and programmes of the public and private sector" there is a specific goal that mentions mining: "Al 2025, el 50% de los sectores productivos considerados relevantes para el cumplimiento de los objetivos del SNAP (minería, silvoagropecuario, energía, obras públicas, transporte, inmobiliario, pesca, acuicultura y turismo) habrán establecido políticas y regulaciones que promuevan prácticas sustentables y coherentes con el SNAP; al 2030, las habrán establecido el 70%" (p. 86). |
| Yes | No | | China | China National Biodiversity | |

| | | | | | |
|---|---|---|---|---|---|
| | | | | Conservation Strategy and Action Plan (2011-2030) | |
| Yes | No | | Colombia | Plan de Accion de Biodiversidad 2016-2030 | There is a recommendation to integrate biodiversity and ecosystem services as fundamental axes in the plans and policies of sectors such as mining (p. 18-19). |
| Yes | Yes | Sand and/or gravel | Comoros | Strategie nationale et plan d'action actualises pour la diversity biologique_v2 | The Objective B1 includes the action B1-5 "Réduire la fiscalité des entreprises de concassage pour limiter l'utilisation des aggrégats marins et appliquer la législation" on the impact of marine sand mining (see the strategy document of 2000). |
| Yes | Yes | both | Congo | Stratégie nationale et plan d'actions sur la diversité biologique (revisé) | Mining of sand and limestone are highlighted as important threats to biodiversity: "un prélèvement de sable lagunaire ayant pour conséquence la perturbation des habitats des espèces, la modification du régime hydrologique des eaux, la perte de la biodiversité benthique" (p. 15, 59). |
| No | No | | Cook Islands | Cook Islands Biodiversity Strategy and Action Plan | |
| Yes | No | | Costa Rica | ESTRATEGIA NACIONAL DE BIODIVERSIDAD 2016-2025 | Within the Strategic Objective 4.4.2.3. "Prevención, protección, seguimiento y control del impacto adverso sobre la biodiversidad y cumplimiento de la legislación ambiental" the text mentions conflicts associated with mines and quarries due to their environmental impacts (p. 50). |
| Yes | No | | Côte d'Ivoire | STRATEGIE ET PLAN D'ACTION POUR LA DIVERSITE BIOLOGIQUE NATIONALE 2016-2020 | The Objective 13 indicates that "By 2020 the exploitation of oil and mineral resources should not compromise the conservation of biological diversity (p. 73). |
| Yes | No | | Croatia | The Nature Protection Strategy and Action Plan of the Republic of | The strategy highlights that "The biggest threat to geodiversity is pressure caused by human activity, in particular excessive exploitation of mineral raw materials, water pollution, interventions on watercourses, illegal waste |

| | | | | | |
|---|---|---|---|---|---|
| | | | | Croatia for the period 2017-2025 | disposal sites, expansion of construction areas, illegal construction and road construction" (p. 8). |
| Yes | No | | Cuba | PROGRAMA NACIONAL SOBRE LA DIVERSIDAD BIOLÓGICA 2016 – 2020 | Several actions and indicators on reducing the environmnetal impacts of mining and increase the surface of restored mines (p. 14, 22). |
| Yes | Yes | Both | Czech Republic | National Biodiversity Strategy of the Czech Republic 2016-2025 | The strategy extensively discuss the theme of mining. It refers to the Raw Materials Policy Strategy (version 6/2016), which imposes the determination of spatial limits and deadlines for mining and quarrying of raw minerals in spatial development policy (p. 75). It mentions that "virtually all stages of the mining process have an impact on the biodiversity", although mined sites (e.g., quarries, sand pits, clay-pits, etc.) can be highly beneficial for biodiversity and nature geodiversity" (p. 76). |
| No | No | | Democratic Republic of Korea | National Biodiversity Stratgey and Action Plan of DPR Korea | |
| Yes | No | | Democratic Republic of the Congo | Stratégie et plan d'action nationaux de la biodiversité 2016-2020. | The strategy highlights issues with pollution associated with the mining of copper and cobalt (p. 46). |
| Yes | Yes | Sand and/or gravel | Denmark | Danish Nature Policy - Our Shared Nature | The dredging for raw materials and "marine construction" are mentioned as a cause of impacts into the marine life (p. 58). |
| Yes | Yes | Sand and/or gravel | Djibouti | STRATÉGIE ET PROGRAMME D’ACTION NATIONAUX POUR LA BIODIVERSITE | Sand and gravel mining are highlighted as a pressure, but the strategy discusses salt mining more specifically as part of the previous biodiversity strategy. It also acknowledges potential impacts on humans: "Ecoulement plus torrentiel, risques de destruction, pertes d’habitats, chute de biodiversité " (p.7). |
| Yes | Yes | Both | Dominica | DOMINICA NATIONAL BIODIVERSITY | The extraction of coastal resources (explicitly sand, gravel, and rocks) threatens coastal biodiversity in Dominica (p. 13). |

| | | | | | |
|---|---|---|---|---|---|
| | | | | STRATEGY AND ACTION PLAN 2014-2020 | |
| Yes | Yes | Sand and/or gravel | Dominican Republic | Estrategia nacional de conservación y uso Sostenible de la Biodiversidad. Plan de accion 2011-2020 | Action 85 refers to the need to strengthen the mechanisms to monitor the extraction and transport of aggregates and the risks at the mining site (p. 95). |
| Yes | No | | Ecuador | ESTRATEGIA NACIONAL DE BIODIVERSIDAD 2015-2030 | The strategy mentions the need of national policies to prevent, control and mitigate the environmental pollution from extractive processes (p. 61). |
| No | No | | Egypt | EGYPTIAN BIODIVERSITY STRATEGY AND ACTION PLAN (2015-2030) | Only the extraction of biological resources is mentioned. |
| Yes | Yes | Both | El Salvador | Estrategia Nacional de Biodiversidad 2013 | The extraction of sand and stones from rivers is highlighted as a cause of soil erosion and sedimentation in wetlands (p. 4, 18). |
| Yes | Yes | Both | Equatorial Guinea | ESTRATEGIA NACIONAL Y PLAN DE ACCIÓN PARA LA CONSERVACIÓN DE LA DIVERSIDAD BIOLÓGICA. ENPADIB | The strategy discusses the regulations and institutions that must regulate the exploitation of quarries and sand resources and propose the reduction of the environmental impacts from mining construction minerals (p. 34). |
| Yes | No | | Eritrea | REVISED NATIONAL BIODIVERSITY STRATEGY AND ACTION PLAN FOR ERITREA (2014-2020) | According to Target 8 "Preventing and mitigating the impacts of pollution and its potential threats the environment will be addressed with a great concern. In view of the current development prospects with an increase in land and marine based activities for agro-industries, mining, port, infrastructure, fishing, livestock, tourism and other sector activities, there is a need for urgent action to prevent and |

| | | | | | |
|---|---|---|---|---|---|
| | | | | | mitigate the impact of the polluting substances, solid and liquid waste that will increasingly be generated across all ecosystems" (p. 60). |
| Yes | Yes | Both | Estonia | Nature Conservation Development Plan until 2020 | The strategy highlights that the main earth resources in Estonia include construction minerals, whose use is regulated by the National Development Plan for the Use of Construction Minerals for 2010–2020: "The problems associated with these earth resources, as well as solutions to the problems, have been scrutinised in the relevant development plans. The most important objective under these plans aims to reduce the negative environmental impact of extraction" (p. 39-40). The Goal 3 "Long-term sustainability of natural resources, and the preconditions for this, are ensured and the principles of the ecosystem approach are followed in the use of natural resources" includes an indicator (Measure 3.2) for "Analysing the impacts of earth resource extraction causing the loss of biodiversity; developing and implementing mitigation measures". |
| Yes | No | | Eswatini | SWAZILAND'S SECOND NATIONAL BIODIVERSITY STRATEGY & ACTION PLAN | Target 11 includes the Action "Develop an Off-setting framework to handle land conversion (e.g. mining)" (p. 38). |
| Yes | No | | Ethiopia | ETHIOPIA'S NATIONAL BIODIVERSITY STRATEGY AND ACTION PLAN 2015-2020 | The strategy highlights that aquatic ecosystems are highly affected by mining (p. 15). |
| Yes | Yes | Sand and/or gravel | Fiji | NATIONAL BIODIVERSITY STRATEGY AND ACTION PLAN FOR FIJI 2020-2025 | Strategic Area SUD4 mentions gravel extraction "Reducing major threats to inland waters (watershed, streams, rivers and lakes such as dredging, floods, gravel extraction, mining, agriculture, deforestation, tourism, sugar, manufacturing, waste management" and Strategic Area SUD5 "Reduce |

| | | | | | |
|---|---|---|---|---|---|
| | | | | | major threats to Fiji's coastal ecosystems such as reclamation, unsustainable tourism development, river dredging and pollution" with associated action plans (p. 46). |
| Yes | Yes | Sand and/or gravel | Finland | Government Resolution on the Strategy for the Conservation and Sustainable Use of Biodiversity in Finland for the years 2012-2020 | The strategy highlights that the environmental risks from commercial marine activities such as gravel extraction are increasing (p. 8) and indicates that "The ecological impacts of projects and plans are difficult to assess with regard to biodiversity values due to the lack of relevant data". |
| No | No | | France | Stratégie nationale pour la biodiversité 2011-2020 | |
| Yes | Yes | Sand and/or gravel | Gabon | Strategie Nationale et plan d'action sur la diversite biologique du Gabon | This strategy highlights coastline erosion has been intensified by human activities including the exploitation of sand on the beaches (p. 51). |
| Yes | Yes | Sand and/or gravel | Gambia | THE NATIONAL BIODIVERSITY STRATEGY AND ACTION PLAN (2015-2020) | Uncontrolled sand/gravel mining are highlighted as anthropogenic drivers of biodiversity loss (p. 15, 42): "In the quest to meet this ever increasing demand, sand mining has become a highly disorganized and chaotic local industry. Although there are attempts by Government to control the activity, illegal sand mining is still common on much of the coastal stretch. Species of marine turtles and water birds habitats have been degraded or totally lost in certain localities. Mining activities along the coast contributes to the process of coastal erosion, threatening many protected ecosystems such as Tanji Bird Reserve, and consequently, the economic and social livelihood of coastal communities" (p. 48). The Target 4 "By 2020, 50% Governments, business and stakeholders have plans for sustainable production and consumption and keep the impacts of resource use within safe ecological limits" mentions sand mining in the technical rationale "Most Government agencies and business sectors have plans but do not reflect biodiversity considerations in |

| | | | | | |
|---|---|---|---|---|---|
| | | | | | their planning and practices therefore, leading to series of environment problems such as logging, sand mining, deforestation etc. Therefore, to adequately inform the planners to mainstream biodiversity issues into their strategies is a key priority for all stakeholders" (p. 63). |
| Yes | No | | Georgia | National Biodiversity Strategy and Action Plan of Georgia 2014-2020 | Within National Target A.3. "By 2020, sustainable use and the economic values of biodiversity and ecosystems are integrated into legislation, national accounting, rural development, agriculture, poverty reduction and other relevant strategies; positive economic incentives have been put in place and incentives harmful to biodiversity have been eliminated or reformed" a Key Objective is to " Integrate biodiversity conservation, sustainable use and ecosystems' values into development programs for such sectors as forestry, energy, agriculture, tourism, mining and infrastructure; take all possible measures to prevent irreversible degradation of ecosystems" (p. 65). |
| No | No | | Germany | Naturschutz-Offensive 2020, für biologische Vielfalt! | |
| Yes | No | | Ghana | NATIONAL BIODIVERISTY STRATEGY AND ACTION PLAN | Illegal surface mining is highlighted as one the greatest threat to biodiversity (p. 11). |
| Yes | Yes | Sand and/or gravel | Greece | National Biodiversity Strategy & Action Plan | Excessive sand extraction due to non-implementation or the partial implementation of the existing institutional framework is highlighted as a cause of biodiversity loss (p. 57). A Specific Target (5.7) aims to ensuring the compatibility of mining activities with biodiversity conservation, including the restoration of quarries (p. 118). |
| Yes | Yes | Sand and/or gravel | Grenada | National Biodiversity Strategy and Action Plan 2016-2020 | Beach sand mining is highlighted as one of the main threats to Grenada's biodiversity in coastal and marine ecosystems (p. 19). For the Strategic Priority 2 "Key National Ecosystems Restored and Sustainably Managed" in the focus area "Coastal and marine biodiversity" a priority action is "Monitor, report and enforce legislation on unsustainable |

| | | | | | |
|---|---|---|---|---|---|
| | | | | | terrestrial practices, coastal infrastructure development, tourism development, mangrove distinction, pollution and waste management, illegal extraction (in particular sand mining) and species, over exploitation and introductions" (p. 34). |
| No | No | | Guatemala | POLÍTICA NACIONAL DE DIVERSIDAD BIOLÓGICA Acuerdo Gubernativo 220-2011 ESTRATEGIA NACIONAL DE DIVERSIDAD BIOLÓGICA Y SU PLAN DE ACCIÓN 2012-2022 Resolución 01-16-2012 del CONAP | |
| Yes | Yes | both | Guinea | STRATÉGIE NATIONALE SUR LA DIVERSITÉ BIOLOGIQUE POUR LA MISE EN ŒUVRE EN GUINÉE DU PLAN STRATEGIQUE 2011 – 2020 ET DES OBJECTIFS D'AICHI | Objective 5 includes an action (5.6) incorporating the restoration plans on impact assessments of mining projects including construction minerals (clay, sand, granite): "Mettre en application les mesures d'accompagnement prévues par les études d'impacts (réhabilitation des sites d'exploitation minière et d'autres carrières : d'argile, de sable, de granite, etc.)" (p. 89). |
| Yes | No | | Guinea-Bissau | Strategy and National Action Plan für Biodiversity 2015-2020 | The strategy discusses the policy framework and environmental implications of the mining sector (p. 89, 95). |
| Yes | No | | Guyana | GUYANA'S NATIONAL BIODIVERSITY STRATEGY AND | Two relevant Priority Actions mention mining: "Promote the integration of biodiversity concerns into mining" (SO2.4) within Target 7 (p. 46), and "To ensure all developers and operators in mining, forestry and agriculture sector are |

| | | | | | |
|---|---|---|---|---|---|
| | | | | ACTION PLAN (2012-2020) | included in the EPA’s environmental authorization process" (SO6.4) within Target 2 (p. 48). |
| Yes | Yes | Sand and/or gravel | Haiti | HAÏTI BIODIVERSITÉ 2030 STRATÉGIE NATIONALE ET PLAN D’ACTIONS POUR LA DIVERSITÉ BIOLOGIQUE (RÉVISÉ-2030) | Dredging for sand and gravel for infrastructure development are highlighted as a cause of negative impacts on biodiversity (p. 109-110). |
| Yes | Yes | Sand and/or gravel | Honduras | ESTRATEGIA NACIONAL DE BIODIVERSIDAD 2018-2022 | Sand and gravel extraction for the construction sector are highlighted as significant human activities associated with urban and tourism expansion (p. 85). |
| Yes | No | | Hungary | National Strategy for the Conservation of Biodiversity in 2015-2020 | Excessive dredging is briefly mentioned as a cause of river degradation (p. 35). |
| Yes | No | | India | NATIONAL BIODIVERISTY ACTION PLAN (NBAP) | |
| Yes | Yes | Limestone | Indonesia | INDONESIAN BIODIVERSITY STRATEGY AND ACTION PLAN 2015-2020 | The conflicts between cement companies (and associated limestone quarries) and conservation in Java (p. 49) and South Sulawesi (p. 130) are discussed and it is indicated that limestone mountains need to be preserved (p. 105). |
| Yes | No | | Iran | Revised National Biodiversity Strategies and Action Plan (NBSAP2) 2016-2030 | Mining is mentioned as an activity that undermines biodiversity (p. 41). |
| Yes | No | | Iraq | IRAQ'S NATIONAL BIODIVERSITY STRATEGY AND | A threat indicator is included on "Mining & Resource Extraction: Studies on oil development, mining & road building impacts, methods and mitigation techniques; |

| | | | | | |
|---|---|---|---|---|---|
| | | | | ACTION PLAN (2015-2020) | Dataset of proposed projects mapped in sensitive areas" (p. 30). It seems that no progress has been made on this indicator. |
| Yes | No | | Ireland | National Biodiversity Action Plan 2017-2021 | Mining and quarrying are recognized as main threats and pressures on Eu protected habitats and species (p. 20). |
| Yes | Yes | Both | Israel | Israel's National Biodiversity Plan | The strategy states that stone quarrying and sand quarrying gradually reduce the size and quality of coastal habitats for these ecosystems' biodiversity (p. 19) |
| Yes | Yes | Both | Jamaica | National Strategy and Action Plan on Biological Diversity in Jamaica 2016-2021 | The strategy describes the "The Quarries Control Act", which concerns to quarry rock, stone, sand, marl, gravel, clay, fill and limestone (p. 25). A gap analysis of the for the sound management of biodiversity in the mining and quarrying sector is presented together with suggestions of mechanisms for improvement (p. 53). Moreover Actions are included for the mainstreaming of biodiversity in the mining and quarrying sector including land-use planning (p. 102), and the inclusion of biodiversity and environmental assessments (p. 107). |
| Yes | Yes | Sand and/or gravel | Japan | The National Biodiversity Strategy of Japan 2012-2020 | The strategy highlights that the disappearance of sandbanks caused by sea gravel extraction in the Seto Inland Sea may have led to a decrease in the population of the Japanese sand lance which is a cornerstone species in the food chain (p. 46). |
| Yes | No | | Jordan | THE NATIONAL BIODIVERSITY STRATEGY AND ACTION PLAN 2015-2020 | Unplanned mining and quarrying are mentioned as a cause of habitat destruction "especially in and around areas of significance to biodiversity such as established and proposed protected areas, important bird areas, genetic resources areas, areas of rangeland diversity and all fragile forest ecosystem" (p. 32). The National Target 12 explicitly mentions mining: "By 2016, renewable energy and mining strategies are reviewed and biodiversity safeguards adopted and enforced" (p. 67). |
| Yes | No | | Kazakhstan | National strategy and action plan on | Mining is highlighted as a sector with environmental impacts on biodiversity (p. 5, 31). |

| | | | | | |
|---|---|---|---|---|---|
| | | | | conservation and sustainable use of biological diversity in the Republic of Kazakhstan | |
| Yes | Yes | Sand and/or gravel | Kiribati | National Biodiversity Strategies and Action Plan 2016-2020 | The management and monitoring of beach mining is included as a national action to address the decline in the turtle nesting beach (p. 26). |
| Yes | No | | Kyrgyzstan | Biodiveristy conservation priorities of the Kyrgyz Republic till 2024 | Mining is recognized as a cause of destruction of natural ecosystems "mining activities carried out in the high, particularly fragile and vulnerable ecosystems, is the source of human disturbance, destruction and pollution of natural ecosystems of habitats of fauna and flora" (p. 4). |
| Yes | No | | Lao PDR | National Biodiversity Strategy and Action Plan 2016-2025 | The mining sector is recognized to be one of the biggest contributors to national revenue in the Lao PDR and have serious environmental implications (p. 21). |
| Yes | Yes | Both | Lebanon | Lebanon's National Biodiversity Strategy and Action Plan | Sand extraction from the shoreline is highlighted as a threat to coastal habitats (p. 23-24) "the high profit from gravels, sands/stones, and the lack of law enforcement are main factors that allow such activities". Quarrying are recognized as drivers of habitat loss and fragmentation "While quarries are needed to support the construction sector, their encroachment on forests and agro-ecosystems is a major problem" "Currently, there are around 1,300 quarries scattered all over the country ranging from small to large scale quarries feeding the cement and construction industries" (p. 23). The Priority Area 5 on Ecosystem Restoration the National Target 9 "By 2030, rehabilitation plans are implemented in at least 20% of degraded sites so that they can safeguard the sustained delivery of ecosystem services" includes the National Action 9.5 "Develop a master plan for the rehabilitation of different types of degraded sites that builds on existing master plans (i.e. quarry and dumpsite rehabilitation)" and National Action 9.7 "Undertake pilot rehabilitation in key sites based on the developed prioritization scheme covering at least one of each type: |

| | | | | | |
|---|---|---|---|---|---|
| | | | | | quarries, dumpsites, degraded forest, rangeland, riverbed, old terraces, and coastal areas" (p. 46). |
| Yes | No | | Lesotho | National Strategy on Lesotho's Biological Diversity: Conservation and Sustainable Use | The Objective 2.4 aims to "Minimize environmental destruction and loss of biodiversity caused by developmental activities" including mining activities. |
| Yes | Yes | Sand and/or gravel | Liberia | NATIONAL BIODIVERSITY STRATEGY AND ACTION PLAN - II 2017-2025 | The strategy highlights that the coastal and marine environment is subjected to erosion due to sand mining (p. 54). |
| No | No | | Liechtenstein | National Biodiversity Strategy and action plan | |
| Yes | No | | Lithuania | ACTION PLAN ON THE CONSERVATION OF LANDSCAPE AND BIOLOGICAL DIVERSITY FOR 2015-2020 | The strategy mentions damaged land by mining operations (p. 7). |
| Yes | No | | Luxembourg | Plan National concernant la Protection de la Nature 2017 - 2021 | The strategy lists former mines as sites of intrest for conservaiton (Kiemerchen / Scheiergronn / Groussebësch, Léiffrächen) (p. 44, 45). |
| Yes | No | | Madagascar | NATIONAL BIODIVERSITY AND ACTION PLANS 2015-2025 | The extractive industries are highlighted as a cause of threat to the biodiversity of Madagascar and the strategy includes the Development of a Strategy of Mining Industry Sector. Within Strategic Objective 4 "In 2025, the Malagasy government and shareholders at all levels will take appropriate steps to implement management plans of resources and maintain the impact of the use of natural resources within limits environmentally safe" there is a key action on "Share best practices on mining exploitation, |

| | | | | | |
|---|---|---|---|---|---|
| | | | | | industrial exploitation, forestry exploitation having a positive impact on forest biodiversity, management of protected areas to promote sustainable production" (p. 98). |
| Yes | No | | Malawi | NATIONAL BIODIVERSITY STRATEGY AND ACTION PLAN II (2015-2025) | Mining is highlighted as a cause of reduction or degradation of important habitats and ecosystems in the country (p. 11). |
| Yes | Yes | Limestone | Malaysia | NATIONAL POLICY ON BIOLOGIVAL DIVERSITY 2016-2025 | The value of limestone (karst) forests is highlighted as a source of marble and raw material for the cement industry and harbouring unique and highgly specialized species (p. 21). Target 7 of the NBSAP on limestone hills protection "By 2025, vulnerable ecosystems and habitats, particularly limestone hills, wetlands, coral reefs and seagrass beds, are adequately protected and restored" (p. 57). |
| Yes | Yes | Sand and/or gravel | Maldives | NBSAP NATIONAL BIODIVERSITY STRATEGY & ACTION PLAN 2016-2025 | The strategy highlights that "While coral and sand mining are controlled through the regulation, it still continues to be one of the core materials in construction and therefore, is exploited" (p. 16) and sand mining is mentioned as cause of biodiversity loss (p. 26). The Target 18 explicitly mentions sand "By 2025, at least 10% of coral reef area, 20% of wetlands and mangroves and at least one sand bank and one uninhabited island from each atoll are under some form of protection and management" (p. 46). |
| Yes | Yes | both | Mali | STRATEGIE NATIONALE ET PLAN D’ACTIONS POUR LA DIVERSITE BIOLOGIQUE, MALI | The strategy inlude actions for the rehabilitation of mines of sand, clay, gold, etc. (p. 87) and discusses financial mechanisms to tax the extraction of aggregates, stone, clay (p. 132). |
| No | No | | Malta | MALTA'S NATIONAL BIODIVERSITY STRATEGY AND ACTION PLAN 2012-2020 | |

| | | | | | |
|---|---|---|---|---|---|
| No | No | | Marshall Islands | The Republic of the Marshall Islands National Biodiversity Strategy and Action Plan | |
| Yes | No | | Mauritania | Stratégie et plan d'action national de la biodiversité 2011-2020 | The impacts of dredging on marine habitats are highlighted (p. 48). |
| Yes | Yes | Sand and/or gravel | Mauritius | National Biodiveristy Strategy and Action Plan 2017-2025 | The strategy highlights the "Removal of Sand Act" as primary legislation on the coastal and marine environment (p. 37). |
| Yes | Yes | Both | Mexico | ESTRATEGIA NACIONAL SOBRE BIODIVERSIDAD DE MEXICO Y PLAN DE ACCION 2016-2030 | Mining of construction minerals are mentioned as a cause of habitat loss and degradation (p. 52) and the strategy includes goals and actions on mining, for example to integrate sustainability criteria in sectoral policies (including those on mining) (specific goal 4.1.5) and to assess and report mining impacts (goal 4.1.3)(p. 118-119). |
| Yes | Yes | Sand and/or gravel | Micronesia | National Biodiveristy Startegy and Action Plan 2018-2023 | Dredging for coastal construction is highlighted as a cause of loss of seagrass beds (p. 12). |
| No | No | | Monaco | Strategie nationale pour la biodiversite Horizon 2030 | |
| Yes | No | | Mongolia | National Biodiversity Programm (2015-2025) | Goal 6 "Protect soil and water resources from chemical and nutrient pollution" includes an objective (13) to "Enable cooperation with government and the general public in the monitoring of legal enforvement of laws regarding chemical pollution from urbanization, mining and manufacturing" (p. 24). |
| Yes | Yes | Sand and/or gravel | Montenegro | NATIONAL BIODIVERSITY STRATEGY WITH THE ACTION PLAN | The strategy mentions construction minerals within target c) "Sustainable agriculture, forestry and water management", with indicator D17 measuring the reduction of the impact of illegal exploitation of gravel and sand (p. 75). |

| | | | | | |
|---|---|---|---|---|---|
| | | | | FOR THE PERIOD 2016-2020 | |
| Yes | Yes | both | Morocco | Stratégie et Plan d'Actions National pour la Diversité Biologique du Maroc, 2016-2020 | The environmental consequences of mining limestone, sand and other resources are highlighted: "Exploitation des mines et carrières : des effondrements ont pour origine l'extraction de calcaires, de gypses, de marnes, de grès et de sables pour les constructions. Aussi, l'exploitation des sols engendre l'appauvrissement des richesses minières. Des paysages défigurés en sont également une conséquence" (p. 22). |
| Yes | No | | Mozambique | NATIONAL STRATEGY AND ACTION PLAN OF BIOLOGICAL DIVERSITY OF MOZAMBIQUE (2015-2035) | The Target 4 "By 2025, define ecologically sustainable systems for the production and consumption based on sustainable practices and adequate investment" includes the Action "4.6. Promote the sustainable use of biodiversity in key productive sectors (mining, agriculture, forests, tourism, energy, public work and housing) according to national and international regulations (e.g. ISO 14001)". Target 7 "Target 7: By 2020, catalog/systematize, disseminate and promote sustainable management practices in agriculture, livestock, aquaculture, mining, forestry and wildlife" includes the action "7.7. Establish and implement sustainable management practices for small-scale mining." |
| Yes | Yes | Limestone | Myanmar | National Biodiversity Strategy and Action Plan 2015-2020 | Limestone quarrying for cement production is highlighted as a majort threat to karst ecosystems (p. 10). Explicit mention of mining in Target 4.1 "By 2020, SEA conducted and guidelines prepared for mining and energy sectors" (p. 45). |
| Yes | No | | Namibia | Namibia'a Second Natinal Biodiversity STRATEGY AND ACTION PLAN 2013-2022 | One of the most critical threats to biodiversity in Namibia is the "Rapid expansion of mining and prospecting" (p. 15). One of the Key Performance Indicators of the Target 7 includes "Compliance with Environmental Management Plans (mining companies)" (p. 23). |
| Yes | Yes | Sand and/or gravel | Nauru | NAURU'S BIODIVERSITY STRATEGY AND ACTION PLAN | The strategy mentions the "Lands Act", which "makes provision for "the leasing of land for ... the removal of trees, crops, soil and sand and the payment of compensation and other moneys" (p. 75). There is a strategy goal to Strategy Goal "To protect Nauru's native biodiversity from impacts of |

| | | | | | |
|---|---|---|---|---|---|
| | | | | | alien invasive species and imported earth materials, through effective border control, effective quarantine and eradication programme" (p. 34). Concerns about phosphate mining are more prominent (p. 69) |
| Yes | Yes | Sand and/or gravel | Nepal | NEPAL NATIONAL BIODIVERSITY STRATEGY AND ACTION PLAN | Widespread gravel mining from streams and river beds is highlighted as a major threat to aquatic biodiversity and cause of deforestation (p. 28,30). A National Target is the "Development and implementation, by 2015, an effective mechanism to control mining of gravel and sand from rivers and streams" (p. 84). |
| Yes | No | | Netherlands | Natural Capital Agenda: conservation and sustainable use of biodiversity | The only mention refers to joining the Coalition of the Willing for a High Seas Marine Protected Area, agreements will be made in international marine management forums (Law of the Sea, fishery, shipping and mineral extraction) for the permanent conservation of the Sargasso Sea. It also suggests that restoration of the shellfish beds in one of the North Sea's protected areas will be done in synergy with activities like sand extraction. |
| Yes | Yes | Sand and/or gravel | New Zealand | Aotearo New Zealand Biodiversity Strategy 2020 | Gravel extraction is highlighted as an activity that can alter or completely destroy habitats (p. 20) and "Commercial catch, seabed and coastal seabed dredging and trawling" is proposed as an indicator for measuring progress towards the strategy outcome 1 "Ecosystems, from mountain tops to ocean depths, are thriving" (p. 70). |
| Yes | No | | Nicaragua | Estrategia Nacional de Biodiversidady su Plan de Acción Nicaragua 2015-2020 | Within goal 7 there is an indicator on incoporating biodiversity conservation criteria in the plans and policies of multiple sectors including mining (p. 53). |
| Yes | Yes | Limestone | Niger | STRATEGIE NATIONALE ET PLAN D'ACTIONS SUR LA DIVERSITE BIOLOGIQUE, 2ème édition | Limestone is highlighted as one of the key minerals extracted in terms of volume (p. 19). |

| | | | | | |
|---|---|---|---|---|---|
| Yes | No | | Nigeria | NATIONAL BIODIVERSITY STRATEGY AND ACTION PLAN 2016-2020 | Uncontrolled, Illegal and Harmful Mining Practices are highlighted as causes of biodiversity loss (p. 19). The National Goal 2 "Reduce the direct pressures on Nigeria's biodiversity resources and promote sustainable use" indicates that "Concerted efforts will be made to promote sustainable practices of land use for agriculture, mining, crude oil exploration, aquaculture, tourism, housing development and industrialization..." (p. 35). |
| Yes | No | | Niue | NIUE NATIONAL BIODIVERSITY STRATEGY AND ACTION PLAN 2015 | Within Objective 8 "Minimise pollution of marine environment" a Key Action (8.1) is to "Ensure that EIAs are conducted for any oil, gas and mineral exploration" (p. 66) and within Objective 2 "Prevent contamination of the Niue groundwater lens" there is a Key Action (2.8) to "Manage mineral extraction to minimise risk of contamination" (p. 82). |
| Yes | Yes | Both | North Macedonia | NATIONAL BIODIVERSITY STRATEGY AND ACTION PLAN For the period 2018-2023 | Open-cast mining (including limestone quarrying) and sand and gravel mining are highlighted as priority threats to biological diversity (p. 58, 74, 162), as these are very often in sensityive areas. |
| Yes | No | | Norway | Norway's national biodiversity action plan | Mining is mentioned as one of the most important pressures to habitats (p. 79). |
| Yes | Yes | Both | Oman | National Biodiversity Strategy and Action Plan | The strategy recognizes that mineral resource exploitation activities, including those for limestone and aggregates may pose threats to natural habitats and wildlife if not addressed properly (p. 29). |
| Yes | Yes | Limestone | Pakistan | National Biodiversity Strategy and Action Plan | The strategy highlights that growing cement industries in the habitat of Punjab Urial in Salt Range pose a threat to the survival of endemic species (p. 58). |
| Yes | Yes | Limestone | Palau | Revised National Biodiversity Strategy and Action Plan 2015-2025 Promoting Wise Development to | Limestone quarrying is highlighted as an immediate threats to partulids (p. 161) and projects such as dredging are considered to be altering habitats in many areas "to such a degree that once abundant marine species are now hard to find" (p. 145). |

| | | | | | |
|---|---|---|---|---|---|
| | | | | Achieve Conservation and Sustainable Use of Biodiversity | |
| Yes | No | | Panama | Plan de Accion Nacional de Biodiversidad 2018-2050 | Mining is highlighted as a key economic sector with influence on biodiversity (p. 55). |
| Yes | No | | Papua New Guinea | National Biodiversity Stragetic Action Plan 2019-2024 | Mining is considered a significant threat to biodiversity as it leads to a large massive habitat loss which affects micro-organisms, vegetation and animals. It mentions gold, copper, nickel, zink, cobalt, and chromite. |
| Yes | No | | Paraguay | EstratEgia NacioNal y PlaN dE acciόN Para la coNsErvacióN dE la BiodivErsidad dEl Paraguay 2015-2020 | Mining is mentioned as a key economic sector (p. 67). |
| Yes | No | | Peru | ESTRATEGIA NACIONAL DE DIVERSIDAD BIOLÓGICA AL 2021 Plan de Acción 2014 - 2018 | Illicit mining is highlighted as a threat to biodiversity in Peru (p. 15). There is a target (70) to complete by 2015 a guide of best practices for biodiversity conservation for mining companies (p. 58). |
| Yes | No | | Philippines | Philippine Biodiveristy Strategy and Action Plan 2015-2028 | Challenges associated with black sand (magnetite) mining are highlighted in the appendix. |
| Yes | Yes | Sand and/or gravel | Poland | The programme of conservation and sustainable use of biodiversity along with Action Plan for the period 2015-2020 | The excessive extraction of sand and gravel is highlighted as a threat to coastal ecosystems (p. 12). |

| | | | | | |
|---|---|---|---|---|---|
| Yes | No | | Portugal | Decreto da Estratégia Nacional de Conservação da Natureza e Biodiversidade 2030 | The strategy includes actions to ensure the conservation of biodiversity and geodiversity in prospecting, research and exploitation of mineral resources (p. 24). |
| Yes | No | | Qatar | Qatar National Biodiversity Strategy and Action Plan 2015-2025 | Dredging and coastal reclamation are highlighted as a cause of stress on the natural marine environment (p. 10). |
| Yes | No | | Republic of Korea | The Republic of Korea's Fourth National Biodiversity Strategy 2019-2023 | Within Strategy 3 "Strengthening biodiversity conservation" and Target 2 "Achieving ecosystem restoration" there is a key action on "launching restoration efforts for damaged ecosystems" including areas damaged by quarries (p. 19). |
| Yes | Yes | Sand and/or gravel | Republic of Moldova | the Strategy on Biological Diversity of the Republic of Moldova for 2015-2020 and the Action Plan for enforcing it | The extraction of sand and gravel is highlighted as a cause of decline of fisheries (p. 22). |
| Yes | Yes | Both | Republic of Sierra Leone | Sierra Leone's Second National Biodiversity Strategy and Action Plan 2017-2026 | Sand mining and dredging are highlighted as causes of degradation/fragmentation/loss of seabed integrity (p. 121). Sand, gravel and rocks (building materials) are amongst the main minerals mined in Sierra Leone and in great demand due to the construction industry and coastal infrastructural development (p. 55). |
| No | No | | Republic of South Sudan | NATIONAL BIODIVERSITY STRATEGY AND ACTION PLAN (2018-2027) | The exploitation of sand and gravel is acknowledged as a contribution of forest biodiversity to the national economy (p. 61). |
| Yes | No | | Romania | National strategy and Action plan for biodiversity conservation 2014-2020 | The "Extension of surface mining activities and extension of areas occupied by wastes without their greening" is listed as a direct threat to Romanian biodiversity (p. 25). |

| | | | | | |
|---|---|---|---|---|---|
| Yes | No | | Russian-Federation | Strategy and Executive Plan for the Conservation of Biodiversity within the Russian Federation | The extraction of minerals is mentioned as a factor responsible for the destruction of Russia's steppe ecosystems (p. 127). |
| Yes | Yes | Both | Rwanda | NATIONAL BIODIVERSITY STRATEGY AND ACTION PLAN | The excessive extraction of boulders, gravel and sand from rivers and streams is highlighted as a direct threat to biodivesity (p. 11, 23), and the production of cement materials exploited in Mashyuza site, is considered by local population as very serious and permanent threat to the survival of biodiversity in these ecosystems and contribute to the disruption of the hydrological cycle and degradation of water quality in streams of the region (p. 23). |
| Yes | Yes | Sand and/or gravel | Saint Kitts and Nevis | ST. CHRISTOPHER (ST. KITTS) & NEVIS NATIONAL BIODIVERSITY STRATEGY & ACTION PLAN 2014-2020 | The strategy highlights that biodiversity (e.g., coral reefs) are under high level of threat from sand mining (p. 18) leading to environmental degradation and habitat loss (p. 19). |
| Yes | Yes | Sand and/or gravel | Saint Lucia | REVISED SECOND NATIONAL BIODIVERSITY STRATEGY AND ACTION PLAN Second NBSAP (2018 - 2025) for SAINT LUCIA | Sand mining is highlighted as an emerging threat to the Leatherback Turtle (Dermochelys coriacea). Specifically the strategy indicates that large-scale sand mining has reduced the nesting habitat of the Grande Anse leatherback beach by more than 50% (p. 64). |
| No | No | | Saint Vincent and The Grenadines | THE NATIONAL BIODIVERSITY STRATEGY AND ACTION PLAN | Sand mining is mentioned in one of the references: US Agency International Development: Caribbean Open Trade Support (USAID-COTS) 2010. Sand mining in St. Vincent and the Grenadines: Impacts and options. |
| Yes | Yes | Sand and/or gravel | Samoa | SAMOA'S NATIONAL BIODIVERSITY STRATEGY AND | The national Target 10 on minimizing pressures from anthropogenic activities on coral reefs, streams and other vulnerable ecosystems (p. 47) includes key actions on conducting a baseline assessment of the coastal sand budget |

| | | | | | |
|---|---|---|---|---|---|
| | | | | ACTION PLAN 2015-2020 | and the enforcement and implementation of planning and permit approval frameworks to reduce coastal reclamation and sand mining activities. |
| No | No | | San Marino | National Biodiversity Strategy and Action Plan | |
| Yes | No | | Sao Tome Principe | NATIONAL BIODIVERSITY STRATEGY AND ACTION PLAN 2015-2020 (NBSAP II) | Mining is mentioned as a threat to corals (p. 36). |
| Yes | Yes | Sand and/or gravel | Saudi Arabia | The National Strategy for Conservation of Biodiversity in the Kingdom of Saudi Arabia | Dredging is highlighted as a threat to marine flora and fauna: "Like landfilling, dredging causes destruction of the marine resources in the dredged area and often has indirect negative impacts from increased sedimentation, which causes a long term destruction of plant and animal communities" (p. 25). "One of the activities most disruptive to coastal and marine resources which causes severe and permanent destruction of coastal habitats such as the loss of mangroves where marine fauna live and reproduce. Also inflicting indirect damage due to sedimentation." (p. 29). Under Strategic Goal 2 (In-situ Conservation - Outside Protected Areas) propose the Action "Limit Landfilling and Dredging Activities on coastal areas". The report also highlights that "The dust resulting from the cement factories at Yanbu has caused health problems for people as well as serious damage to nearby coral reefs and green turtle nesting sites." (p. 28) Further research reveals that impacts are due to heavy metals released from the cement factory but limestone mining is not mentioned. |
| Yes | Yes | Sand and/or gravel | Senegal | Stratégie nationale & plan national d'actions pour la biodiversité | The extraction of marine sand is highlighted as a threat coastal and marine ecosystems (p. 31). |
| Yes | Yes | Sand and/or gravel | Serbia | BIODIVERSITY STRATEGY OF THE REPUBLIC OF | The extraction of gravel and alluvial sands and dredging are highlighted as factors that alter flow regimes in natural waterways (p. 55). |

| | | | | | |
|---|---|---|---|---|---|
| | | | | SERBIOA FOR THE PERIOD 2011-2018 | |
| Yes | No | | Seychelles | Seychelle's National Biodiversity Strategy and Action Plan 2015-2020 | Dredging is listed among the causes of decline of seagrass (p. 36). |
| No | No | | Singapore | CONSERVING OUR BIODIVERSITY Singapore's National Bodiversity Strategy and Action Plan | The Ketam Mountain Bike Park in Pulau Ubin is shown as a good example of rehabilitation. The site had been highly impacted by past granite quarrying activities (p. 9). |
| Yes | Yes | Sand and/or gravel | Slovakia | Updated National Strategy for the Protection of Biodiversity to 2020 | The National Target C.6 includes ensuring adequate protection for aquatic and water dependent habitats requires to prevent the unjustified gravel extraction from river beds (p. 20). |
| Yes | Yes | Both | Slovenia | National Environmental Action Programme 2020–2030 | The strategy highlights that abandoned gravel pits and karst caves are a particular problem. Several mentions to the need of material efficiency strategies for raw materials (p. 67). |
| Yes | No | | Solomon Islands | THE NATIONAL BIODIVERSITY STRATGIC ACTION PLAN 2016-2020 | The strategy indicates that inland water systems and coastal systems are under threat from mining (p. 32). The Mines and Mineral Act provides the provision for integrating conservation with mining development for which the latest NBSAP "provides the necessary road map for implementing the Act and in particular relevant to management of waste and, pollution control as popularly associated with mining development. The corresponding targets and action points, are therefore set the roadmap for implementing the Act within the scope of biodiversity." (p. 50) |
| Yes | Yes | Sand and/or gravel | Somalia | National Biodiversity Strategy and Action Plan (NBSAP) | Sand mining is listed as a driver of change for the coastal biodiversity (p. 53). |
| Yes | No | | South Africa | South Africa's 2nd National Biodiversity | Within Target 3.2 "Embed biodiversity considerations into national, provincial and municipal development planning and monitoring" there is an action planned (3.2.7.) to integrate |

| | | | | | |
|---|---|---|---|---|---|
| | | | | Strategy and Action Plan 2015-2025 | biodiversity priorities into key production sector strategies and plans, including for agriculture, mariculture, aquaculture, mining, forestry, water, land reform and rural development, through cooperative approaches (p. 42). Within Target 3.3 "Strengthen and streamline development authorisations and decision-making" there is an indicator on "Number of environmentally significant areas identified and published for restriction for mining activities" (p. 43). Within Target 6.2 "The status of species and ecosystems is regularly monitored and assessed" there is an action planned (6.2.5) on "Regularly map key pressures on biodiversity, including landcover change, pressures in the marine environments, such as fisheries, trawling, mining, and the density and distribution of invasive alien species". |
| Yes | Yes | Both | Spain | Plan estratégico del patrimonio natural y de la biodiversidad 2011-2017 | Aggregate extraction is highlighted as a possible cause of geomorphological changes with severe impacts on river and coastal ecosystems (p. 71). |
| Yes | Yes | Sand and/or gravel | Sri Lanka | National Biodiversity Strategic Action Plan 2016-2022 | River and beach sand mining are listed as causes of habitat degradation (p. 63). A National Policy on Sand as a Resource for the Construction Industry from 2006 is mentioned among other relevant policies (p. 86). |
| Yes | No | | Sudan | National Biodiversity Strategy and Action Plan 2015-2020 | To tackle Aichi Target 8 "By 2020, pollution, including from excess nutrients, has been brought to levels that are not detrimental to ecosystem function and biodiversity" one of the Actions proposed is to "Reduce to environmentally acceptable levels the adverse impacts of tradition as well as organized gold mining on wildlife and inland waters and marine habitat" (p. 68). |
| Yes | No | | Suriname | National Biodiversity Action Plan (NBAP) 2012-2016 | Within the Objective 1 "Conservation of biodiversity" the Sub-objective 1.4 indicates that "Responsible mining with minimisation of damage to the environment and biodiversity and environmental restoration" (p. 27). |
| Yes | Yes | Sand and/or gravel | Sweden | A Strategy for Biodiversity and Ecosystem Services & | The accessible pdf document of the National Biodiversity Strategy and Action Plan (v.3) submitted on 2016-06-30 shows only a a translation of relevant parts of Government |

| | | | | | |
|---|---|---|---|---|---|
| | | | | Action Plan on Biological Diversity | bill on biodiversity and ecosystem services A Swedish strategy for biodiversity and ecosystem services Gov. Bill 2013/14:141 (21 pages). Given that this is not comprehensive, we also checked the previous National Biodiversity Strategy and Action Plan (v.2 submitted on 2007). The NBSAP v.2 highlights the extraction of sand and gravel from coastal areas as a threat to biodiversity "The use of coastal areas for activities such as fishing, aquaculture, engineering projects (marinas, roads, bridges, shipping lanes, dredging), recreation (tourist centres, summer cottages) and the extraction of mineral resources (oil and gas, sand) poses threats to both species and habitats. The trend today is one of increasing pressure on Swedish coastal and sea areas, and hence a growing threat to biological diversity. The combined effect of many minor encroachments can be considerable." (p. 66). |
| Yes | No | | Switzerland | Swiss Biodiversity Strategy | Raw materials extraction, production, use, disposal and the recycling of these goods are recognized as causes of direct or indirect impacts on global biodiversity" (p. 19) and highlights that the on-going biodiversity loss harbours various corporate risks for businesses including mining, such as shortages of and increases in the cost of raw materials (p. 43). |
| No | No | | Thailand | Master Plan for Integrated Biodiversity Management | |
| Yes | Yes | Sand and/or gravel | Timor-Leste | The National Biodiversity Strategy and Action Plan of Timor-Leste (2011-2020) | The extraction of sand and stones in riverbeds is highlighted as a cause of pollution and sedimentation in freshwater and inland ecosystems (p. 6). |
| Yes | Yes | both | Togo | Stratégie et Plan d’Action National pour la Biodiversité du Togo SPANB 2011-2020 | Sand and limestone mining are highlighted as relevant activities to consider in land-use planning: "les établissements constitués des différentes agglomérations (villes, villages), les infrastructures, les terres servant aux extractions minières (carrière d’extraction de calcaire, de |

| | | | | | |
|---|---|---|---|---|---|
| | | | | | phosphate, gneiss, marbre) et les sols nus (59 000 ha)." (p. 22) |
| Yes | Yes | Sand and/or gravel | Tonga | National Biodiversity Strategy & Action Plan | Mining threats to forests are highlighted: "Others more exposed to threats, particularly mangroves, needs protection from reclamation, unlawful mining activities". In Theme Area 2 "Marine Ecosystems Basis for Action" the NBSAP highlights that "the ability of coastal and marine ecosystems to perform these functions have been severely impaired by poorly planned development activities including infrastructure, land reclamation, sand mining, waste disposal and settlements". The NBSAP includes as key action under Objective 2.1 "Managing Impacts of Land-based activities" to “Strengthen existing legislation and or introduce new ones to support effective EIA procedures as a means of regulating sand mining, land reclamation, coral quarrying, mangrove destruction and waste disposal” (p. 28). |
| Yes | No | | Trinidad and Tobago | BIODIVERSITY STRATEGY & ACTION PLAN FOR TRINIDAD AND TOBAGO 2017-2022 | One of the indicators for Monitoring and Evaluation of the NBSAP is "Number of illegal bush fires and quarrying" to be measured anually (p. 116). |
| No | No | | Tunisia | NATIONALE ET DU PLAN D’ACTION NATIONAUX SUR LA BIODIVERSITE<br><br>Stratégie et plan d’action nationaux pour la biodiversité 2018-2030 | The strategy discusses "ré-ensablement" as a method to protect and restore degraded coastal ecosystems "La protection et la restauration des côtes les plus dégradées/menacées moyennant (i) la fixation du trait de côte, (ii) le ré-ensablement, (iii) la mise en place d’éco-plages, etc." (p. 52). |
| No | No | | Turkey | NATIONAL BIODIVERISTY ACTION PLAN 2018-2028 | |
| Yes | Yes | Both | Tuvalu | Tuvalu National Biodiversity Strategy | The strategy mentions the Foreshore and Land Reclamation Act: "the removal of sand, gravel, reef mud, coral, rock and |

| | | | | | |
|---|---|---|---|---|---|
| | | | | and Action Plan 2012-2016 | other may only be taken from foreshore areas with the approval of the relevant island council" (p. 66). |
| Yes | Yes | Sand and/or gravel | Uganda | NATIONAL BIODIVERSITY STRATEGY AND ACTION PLAN II (2015-2025) | The harvesting of construction materials like clay and sand is highlighted as a current threat to wetlands and their biodiversity (p. 13). |
| Yes | Yes | Limestone | Ukraine | THE MAIN PRINCIPLES (STRATEGY) OF THE NATIONAL ENVIRONMENTAL POLICY OF UKRAINE UNTIL 2020 | Limestone and other construction stone dominate the volume of production of minerals in Ukraine. The NBSAP indicate that intensive mining (in general) "has resulted in considerable changes to the geological environment and occurrence of natural and technogenic emergencies (p. 2-3). |
| No | No | | United Kingdom | Biodiversity 2020: A strategy for England's wildlife and ecosystem services | |
| Yes | Yes | Sand and/or gravel | United Republic of Tanzania | National Biodiversity Strategy and Action Plan (NBSAP) 2015-2020 | Mining of minerals and aggregate is highlighted as serious threats to habitats (p. 41). |
| Yes | No | | Uruguay | Estrategia Nacional para la Conservación y Uso Sostenible de la Diversidad Biológica del Uruguay 2016-2020 | The strategy includes an action to contribute to the process of developing guidance and best practices for mining activities (p. 47). |
| Yes | Yes | Sand and/or gravel | Vanuatu | VANUATU NATIONAL BIODIVERSITY STRATEGY AND ACTION PLAN (NSAP) 2018-2030 | The strategy includes 'stopping marine sand extraction by 2020' as a key action to address the decline of marine resources (p. 154). |

| | | | | | |
|---|---|---|---|---|---|
| Yes | No | | Venezuela | Estrategia Nacional para la Conservación de la Diversidad Biológica 2010-2020 y su Plan de Acción Nacional | The strategy includes an Action to prioritize large-scale mining (p. 52). |
| Yes | Yes | Both | Viet Nam | VIETNAM NATIONAL BIODIVERSITY STRATEGY TO 2020, VISION TO 2030 | Quarrying of limestone and overexploitation of river sand and gravel are highlighted as direct causes of biodiversity degradation in the country (p. 46). |
| Yes | Yes | Sand and/or gravel | Yemen | National Biodiversity Strategy and Action Plan II "achieving a resilient, productive and sustainable socio-ecosystem by 2050" | The extraction of minerals is highlighted as a cause of widepsread ecosystem loss. There are considerations on dredging and quarrying on coastal habitats: "Habitats are fast deteriorating to meet increased land reclamation for coastal urbanization, industrial growth, oil exploration, fishing; tourism; agriculture; aquaculture, sea water desalination and ports & sewage development. These activities are broadly associated with extensive dredging, land filling, mining and quarrying with subsequent loss of the Red Sea & Arabian sea coastal habitats such as coral reef, mangrove, wetland, palm trees, lagoons, beaches (sandy & rocky), dunes, Sabkha, Seagrass Beds & Turtle Nesting Sites"(p. 36-37). |
| Yes | No | | Zambia | ZAMBIA'S SECOND NATIONAL BIODIVERSITY STRATEGY AND ACTION PLAN (NBSAP -2) 2015-2020 | Mining is highlighted as a cause of habitat transformation and biodiversity loss y several protected areas (p. 23). The strategy includes a Strategic Intervention to "Oblige the mining industry to contribute to the Environmental Protection Fund (EPF) under the Mines and Minerals Development Act" (p. 56). |
| Yes | Yes | Sand and/or gravel | Zimbabwe | NATIONAL BIODIVERSITY STRATEGY AND ACTION PLAN 2014 | River sand mining is highlighted as a threat to aquatic habitats and associated biodiversity (p. 13). |
