## Appendix S4 for "Mining threats in high-level biodiversity conservation policies"

**Appendix S4**. Detailed search results on IPBES reports. In bold red the search term identified in the report's quote. Detailed search results on IPBES assessments. Candidate assessments were: Global, Land Degradation and Restoration, Asia-Pacific, Europe and Central Asia, Americas, Africa. In bold red the search term identified in the assessment's quote. Quotes were selected based on their alignment with the subject of mining construction minerals.

| Assessment | Chapter | Term | Selected quotes |
|---|---|---|---|
| Global | Status and trends – Drivers of change | dredg | *"Large-scale **dredging** has occurred in several countries in Asia and the Middle East, beyond the near-shore environments, for creation of airports, tourism facilities and islands. Land reclamation is linked to the degradation of wetlands, seagrass beds and decreased coastal water quality, with negative impacts on regional groundwater regimes discharges to the coasts."* |
| | | mining | *Section 2.1.11.5 **Mining*** |
| | | mining | *"Coastal areas intensively and multiply used by humans is an anthrome, defined by artificial constructions linked to human settlements, industry, aquaculture, or infrastructure that transforms coastal habitats (Baueret al. 2015). These include a) coastal defences (breakwaters, groynes, and jetties), b) coastal protection (seawalls, bulkheads, and pilings), c) floating docks, e) artificial islands, f) dumping and **mining** areas, g) artificial structures for energy (including renewable energies) and h) port development and coastal support"* |
| | | sand, gravel | *"Changes in the water inflows and abstraction, and structural modifications (e.g. drainage and conversion) all directly drive the loss of inland wetlands (Ramsar 2018). Indirect drivers include overfishing, intensive wood harvesting (e.g. in wetland forests), peat extraction, and **sand** and **gravel** extraction for construction (Ramsar 2018)"* |
| Land degradation | Direct and indirect drivers of land degradation and restoration | cement | *"Anthropogenic activities including fossil fuel combustion, **cement** production, deforestation and land-use change have resulted in significant amounts of greenhouse gases (GHGs) being emitted into the atmosphere, and atmospheric concentrations of GHGs including carbon dioxide (CO2), methane (CH4) and nitrous oxide (N2O) are unprecedented in at least the last 800,000 years (IPCC, 2014a)"* |
| | | mineral, sand | *"The spatial scales over which different direct anthropogenic drivers are manifested ranges from local (e.g., land conversion or localized **mineral** or **sand** extraction) to regional (e.g., invasive species) or global scales (e.g., climate change)."* |
| | | mining, mineral | *"Geographic databases compiled by the United States Geological Survey (USGS) (Matos et al., 2015) show more than 17,000 different large-scale **mining** sites in 171 countries. Hundreds of different **mineral** commodities are mined for diverse uses including energy generation, construction, manufacturing and industry, fertilizers, electronics, and medicine. As negative impacts from extractive industries expand and more directly harm ecosystem services and biodiversity, countries have begun to regulate or incentivize restoration of abandoned mine lands as a way to recover some of the ecosystem services lost* |

| | | | |
|---|---|---|---|
| | | | *during the extractive process (Bradshaw, 1997; Cooke & Johnson, 2002; Bridge, 2004)"* |
| | | mineral, concrete, mining, gravel | *"Extractive industries can be usefully broken down into six distinct categories… These industries are: (1) ferrous **minerals** (iron, nickel, titanium and so on); (2) non-ferrous **minerals** (copper, gallium, aluminum); (3) liquid and gaseous fuels (oil and gas); (4) **mineral** fuels (coal and uranium); (5) industrial **minerals** (salt, **concrete**, gypsum, phosphate); and (6) precious metals (silver, gold, platinum). These industries are broadly associated with one of three extraction techniques that are highly variable in their disturbance and subsequent impacts on the land surface: (i) underground **mining**, where a shaft is dug into the earth and **minerals** are brought to the surface resulting in relatively small spatial surface impact directly from **mining**; (ii) surface and open pit **mining** with resource seams directly accessed from the surface with a much larger footprint of surface disturbance (coal, iron, copper, lithium, **gravel**, phosphates); and (iii) well extraction where a small platform is built to hold a well for extracting liquid and gaseous resources (gas and oil)."* |
| Land degradation | Status and trends of land degradation and restoration and associated changes in biodiversity and ecosystem functions | cement | *"Primary human mercury emissions include artisanal gold mining (37%), coal combustion (24%), non-ferrous metal production (10%) and **cement** production (9%) (UNEP, 2013)."* |
| | | dredg | *"In many areas, excavation and earth-moving have changed flood patterns, created barriers to runoff and erosion, funnelled sediments into new deposition areas, created unstable spoil heaps, and **dredged** sediment from water bodies to create new land with consequent starving of existing beaches"* |
| | | aggregates, sand, gravel, concrete | *"In contrast, construction **aggregates** like **sand** and **gravel** used in **concrete**, asphalt and building materials are low value per weight so sourcing aggregates is often done more locally with the consequent environmental impacts more globally dispersed (Langer, 2009)"* |
| Asia Pacific | Status, trends and future dynamics of Biodiversity and ecosystems underpinning nature's contributions to people | cement | *"Terrestrial ecosystems that are distinct from the regional type expected for that particular climate, as a result of unusual and extreme geology and/or soils, can make a major contribution to the regional diversity of plants and animals. Whereas these ecosystems have often been treated as wasteland and given no protection, those are now under rapidly increasing threats due to the demand for **cement** and other products."* |
| | | limestone, quarr, cement | *"In SE Asia, **limestone** karsts are often found in areas near development and support remnants of ecosystems which previously had wider distributions but have since been lost to development. The major threat to the survival of karst-associated species is **quarrying** (Sodhi & Brook, 2006). A conservative figure of globally threatened karst-associated species listed by IUCN as critically endangered, endangered, or vulnerable stood at 143 species and of these 31 species (ca. 21 per cent) occur in South-East Asia (Clements et al., 2006). With good financial returns from karst **quarrying** for **cement** manufacturing, it is unlikely this* |

| | | |
|---|---|---|
| | | *exploitation will be slowed down or halted, more so in some SE Asia countries where karst protection is minimal or non-existent (e.g., Myanmar, Cambodia). Current laws for the protection of* ***limestone*** *karst in several countries in the Asia-Pacific region, if any, are lacking, lax and ineffective (Kiew, 2001; Lim & Cranbrook, 2002). An example is the case of Malaysia, where majority of the* ***limestone*** *hills are classified as State Forest Land and do not have protected area status hence vulnerable to anthropogenic disturbances (Clements et al.,2006; Liew et al., 2016).”* |
| | sand, gravel, mining | *“However, one of the important and distinctive landforms that remains least documented along coastal areas of the Asia-Pacific region is ‘Beaches and Rocky Shores’. They include rick shingle beaches and* ***sand****bars, rocky headlands and cliffs along subtidal and intertidal habitats. These habitats are reported to be more threatened due to* ***sand*** *and* ***gravel mining*** *compared to others (Butler & Bax, 2014; Peduzzi, 2014; Thaman, 2013; UNEP/UNCTAD, 2014).”* |
| Direct and indirect drivers of change in biodiversity and nature’s contributions to people | mineral | *“As the rapid urbanization and industrialization in the Asia-Pacific economies, the use of primary materials (metal ores and industrial* ***minerals****, fossil fuels and construction* ***minerals****) continues to grow (Fong-Sam et al., 2016; UNEP, 2016a; 2016b)”* |
| | dredg, mining, sand, gravel | *“Human induced activities such as bottom trawling (dragging fishing nets along the seafloor), harbour* ***dredging****, break water constructions (for the development of ports and harbours),* ***mining*** *(seabed/****sand mining****,* ***gravel*** *extraction and other extractive industries) and growing amounts of marine pollution (including plastic waste) destroy critical marine habitats, physical damage to the ocean floor, accelerate sea erosion or coastline changes (Dattatri, 2015; United Nations, 1992)”* (p. 322) |
| | limestone, cement, quarr, mineral | *“Rapid economic growth and expansion of transportation and utility infrastructure has seen the demand for* ***cement*** *in Asia-Pacific countries increased simultaneously. Currently, India, Vietnam, and Malaysia are among the top five exporters of* ***limestone*** *in the world and export near 20 per cent of global* ***cement*** *collectively (A. C. Hughes, 2017)”* (p. 276) …*“In the Asia-Pacific region high dependence on* ***mineral*** *extraction resulted in vegetation clearance and ecosystem degradation through access expanding, exploration drilling, overburden stripping, ground-water pollution, and* ***quarry*** *collapse. Indirect effects include fragmentation of landscape through road and settlement construction or threats to human health due to air pollution such as dusts or smelter emissions (International Council on Mining and Metals (ICMM), 2006; Y. Y. Yang et al., 2014). The extensive exploitation of* ***limestone*** *for* ***cement*** *production has had devastating consequences on ecosystems in karst areas which represent a few global endemicity hotspots (A. C. Hughes, 2017). The depletion of* ***mineral*** *resources is increasingly being proposed in remote and biodiversity-rich areas that were previously unexplored (International Council on Mining and Metals (ICMM), 2006) or even protected (Durán et al., 2013)”* (p. 276) |

| | | | |
|---|---|---|---|
| Europe Central Asia | Status, trends and future dynamics of Biodiversity and ecosystems underpinning nature's contributions to people | gravel, dredg | *"Modification of rivers by straightening channels, **dredging** and canalizing, has resulted in the loss of species that inhabit or breed in **gravel** beds and river margins."; "According to a national reports review undertaken by the Secretariat of the Convention (Ramsar, 2015a, 2015b), Ramsar wetlands in the region face increasing pressures from rapid urbanization and land-use changes for tourism, infrastructure development (transport and energy) and non-sustainable exploitation of natural resources (e.g. water, **gravel**, peat, oil, gas)"* |
| | Direct and indirect drivers of change in biodiversity and nature's contributions to people | gravel, sand | *"Fundamentally, economic growth is largely explained by investments in real capital and there is a near-linear relationship between GDP growth and physical capital accumulation in most countries (Malmaeus, 2016). There are, in turn, clear correlations between investments in physical capital, and resource use including metals (Chen & Graedel, 2015; Kondo et al., 2012), **gravel** and **sand** (UNEP, 2014), and biomass."* |
| | | mineral, dredg | *"In Western and Central Europe, extraction of abiotic resources is highly dominated by construction and industrial **minerals**, and to a more limited extent fossil energy (Bahn-Walkowiak et al., 2012). **Dredging** and pumping operations have a direct effect on the local biological communities and cause changes in the composition of fauna (Pérez-Ruzafa et al., 2007), and reduction in species diversity, abundance, and biomass (Bolam et al., 2015; Sutton et al., 2009). This changes food webs, particularly lower trophic levels including detritivores, with impacts on carbon cycling (Tecchio et al., 2016)."* |
| | | mineral | *"In Eastern Europe and Central Asia during the Soviet era, agricultural enterprises fulfilled many obligations related to providing jobs and services to local population (e.g., schools, shops, centres of culture, libraries etc.) (Figure 4.14). Since the 1990s these obligations have been transferred to local governments, which have not had resources to fulfil them (Ioffe et al., 2012). This led to a sharp increase in the burden on the biological resources of rural areas (e.g. through poaching and illegal logging); destructive extraction of soil and **mineral** resources (e.g. through sale of fertile topsoil and illegal mass extraction of building materials and coal); as well as growing poverty in rural areas (Allina-Pisano, 2007; Ovcharova & Pishnyak, 2003; Petrick et al., 2013; Visser & Schoenmaker, 2011)"* |
| Europe Central Asia | Options for governance and decision-making across scales and sectors | cement | *"There is a widely recognized gap in existing regulations on spatial planning of industrial activities, especially in a transboundary context. Often industrial facilities (such as hydropower plants, **cement** plants or coal mines) are placed at inappropriate locations where they cause huge damage to biodiversity and ecosystem services. Strategic assessments of sectoral development schemes and programmes are often employed to direct development away from sensitive areas."* |
| Americas | ; Status, trends and future dynamics of Biodiversity and ecosystems | dredg | *"Many wetlands in urban areas that have been modified by filling or **dredging** experience high pulses of stormwater from watersheds with diminished infiltration, and receive toxins from transportation (e.g. chloride from road de-icing* |

| | | | |
|---|---|---|---|
| | underpinning nature's contributions to people | | *salts) and industrial run-off (Brinson & Malvárez, 2002; Sanzo & Hecnar, 2006; Federal Provincial Territorial Governments of Canada, 2010)."* |
| | Direct and indirect drivers of change in biodiversity and nature's contributions to people | mining, cement | *"Major sources of atmospheric mercury include fossil fuel (primarily coal) combustion (the largest source), artisanal gold* ***mining****, non-ferrous metal manufacturing,* ***cement*** *production, waste disposal, caustic soda production, and emissions from soils, sediment, water, and biomass burning, including re-emissions from past anthropogenic emissions (Pacnya et al.,2006; Pirrone et al., 2010)."* |
| | | mining | *"Pollution from past and ongoing coal* ***mining****, hard-rock* ***mining****, and metal-ore smelting, expose humans, fish and wildlife to toxicants (e.g. toxic metals and selenium) across North America; thousands of mines are abandoned, and bankrupcies of mining companies are common, leaving neither public nor private funds available to to mitigate or restore these sites and allowing toxic releases and exposures to continue"* |
| | Current and Future Interactions between Nature and Society | dredg | *"Although seagrass declines have been related to a combination of impacts rather than individual threats (Orth et al., 2006), two major causes of loss were identified by Waycott et al., 2009: direct impacts from coastal development and* ***dredging*** *activities"* |
| Africa | Status, trends and future dynamics of Biodiversity and ecosystems underpinning nature's contributions to people | sand, mining | *"Marine resources include commercially valuable fish that are exploited at artisanal and industrial scales. The exploitable species of aquatic fauna within the marine and coastal ecosystems consist essentially of fishes, shrimps and molluscs. Currently the Carangidae, Carcharinidae, Clupeidae, Elopidae, Ephippidae, Haemulidae, Lutjanidae, Paralichthyidae, Polynemidae, Mugilidae, Sciaenidae families are overexploited (Ogandagas, 2003). An accelerated growth of coastal populations has led to crowded conditions where the poor depend on subsistence activities such as fishing, farming,* ***sand*** *and salt* ***mining*** *and production of charcoal (Sherman et al.,2008)."; "In West Africa, mangroves are found discontinuously from Senegal to the Niger Delta, however, these mangroves are in moderate decline, with an estimated average decline of 25% between 1980 and 2006, then recovering in a few countries in the last decade. The decline is due to cutting of the trees for fuelwood and poles for housing construction; urbanisation and industrialisation; the use of poison and dynamite for fishing, canalisation, discharge of sewage and other pollutants, siltation,* ***sand mining****, erosion, construction of embankments; and in some areas, from the damming of the Volta River."* |
| | Direct and indirect drivers of change in biodiversity and nature's contributions to people | mining, dredg | *"They [community-based conservation initiatives] may also suffer of changing value systems, increased pressure on natural resources and other internal tensions. They are exposed to both external and internal threats: imposed development and resource exploitation processes, such as* ***mining*** *and resource extraction, logging, tree plantation, industrial fishing, sea* ***dredging****, land conversion to large-scale grazing or agriculture, urbanisation and major infrastructure (roads, ports, airports, dams,tourism)."* |
