## Appendix S5 for "Mining threats in high-level biodiversity conservation policies"

**Appendix S5**. Multi-lateral Environmental Agreements (MEAs) that are relevant to biodiversity conservation that include direct or indirect reference to the mining of construction minerals. This list does not include regional conventions that directly or indirectly refer to aggregate exploitation nor reports nor documents providing guidance and/or standards.

| MEA | Focus | Year | Relevance for construction minerals | Source / Link |
|---|---|---|---|---|
| Ramsar convention | Wetlands | 1971 | The Ramsar Convention on Wetlands of International Importance provides an international framework for “the conservation and wise use of wetlands and their resources”. Follow-up documents such as the Global wetland outlook (Gardner and Finlayson, 2018) have identified the extraction of sand and gravel from rivers and coasts as a direct driver of change in wetlands. | https://www.ramsar.org/about/the-convention-on-wetlands-and-its-mission |
| WHC | Human activities | 1972 | The World Heritage Convention (WHC) is an international treaty signed on 23 November 1972, which created the World Heritage Sites, with the primary goals of protecting cultural and natural heritage of outstanding universal value. The international status of a World Heritage site should provide protection against mineral exploration and exploitation. | https://whc.unesco.org/en/conventiontext/ |
| UNCLOS | Seabed aggregates | 1982 | United Nations convention on the Law of the Sea (UNCLOS) adopted in 1982 (ratified by 168 parties) aimed to ensure that no significant harm to biodiversity in marine environments is caused from activities including aggregates mining. It provides for the delimitation of maritime zones and regulates rights and obligations in respect of usage, development and preservation for these zones, including resource mining. For instance, its rules devoted to the preservation of the marine environment have been applied to sand mining by the Permanent Court of Arbitration in the South China Sea Arbitration and Malaysia against Singapore concerning land reclamation and dredging in and around the Straits of Johor. However, experts highlight that “as no direct international standards for marine aggregate extraction are in place, this provision falls back on applying the precautionary approach and employing best environmental practice, with their effectiveness and interpretation being tied to each member country’s regulatory efforts” (Peduzzi, 2014; Weyman, 2016). | https://www.un.org/depts/los/convention_agreements/texts/unclos/unclos_e.pdf<br>https://pcacases.com/web/sendAttach/2086<br>https://www.itlos.org/fileadmin/itlos/documents/cases/case_no_12/12_order_081003_en.pdf |
| CBD and Río Declaration | Natural resources, human pressures | 1992 | The Convention on Biological Diversity (CBD) was adopted on 1992 and was formally enforced on the 29th of December, 1993. The CBD aimed at encouraging cooperation among countries for the “conservation of biological diversity, the sustainable use of its components and the fair and equitable sharing of the benefits…”. The Convention currently has 193 parties. According to Art. 3 of the CBD, the States could still retain and pursue the right to exploit their resources but will b obligated to do | https://www.cbd.int/doc/legal/cbd-en.pdf<br>https://www.un.org/en/development/desa/population/migration/generalassembly/docs/globalcompact/A_CO |

| | | | | |
|---|---|---|---|---|
| | | | so responsibly and ensure that they do not damage the environment within their jurisdiction, or in the jurisdiction of other States. The CBD exists as a legal framework for all the member countries. Arts. 5, 6 and 7 encourage cooperation among States to develop conservation strategies and programmes for ensuring sustainable use of natural resources, while simultaneously identifying, monitoring, and rehabilitating those environments which have been drastically damaged. Art. 7, in particular, imposes a duty on the member States to identify, monitor and assess the impact of activities like sand mining on the environment. Art. 14 of the Convention places an obligation on the member States to also carry out assessments to determine the environmental impact of proposed infrastructure projects. In 1992, the UN Conference on Environment and Development adopted the Río Declaration on the Environment and Development, which proclaims in Principle 15 the precautionary approach. | NF.151_26_Vol.I_Declaration.pdf |
| UNCCD | Mining as a general pressure | 1994 | The UN Convention to Combat Desertification (UNCCD) was adopted in 1994. Two years since the Convention entered into force on 26 December 1996. The objective of the UNCCD is to combat desertification and land degradation, and to mitigate the effects of drought in affected countries around the world, particularly in Africa, through effective action at all levels. Several reports deriving from this convention, such as the Global Land Outlook include sand mining as an important pressure to consider, national reports (e.g., Samoa; Bahamas) and reviews for the implementation of the convention include reports of illegal sand mining that concern the implementation of the goals of the UNCCD (e.g., Saint Kitts and Nevis; Antigua y Barbuda). | https://www.sprep.org/att/IRC/eCOPIES/Countries/Samoa/40.pdf<br>https://www.unccd.int/sites/default/files/naps/bahamas-eng2006.pdf<br>ICCD/COP(4)/AHWG/3/Add.2<br>ICCD/CRIC(1)/4/Add.2 |
| Protocol of the London Convention | Seabed aggregates | 1996 | Protocol of the 1972 Convention on the Prevention of Marine Pollution by Dumping of Wastes and Other Matter (London Convention). The London Convention was crafted to establish a global commitment to regulate all sources of pollution in the marine environment, emphasizing the prevention of polluting the sea by dumping matter that creates hazards or damage the marine environment. However, dredged material and any "matter directly arising from, or related to the exploration, exploitation and associated off-shore processing of sea-bed mineral resources" are explicitly exempt from its definition of dumping. To strengthen the regulations under the initial convention, the contracting parties agreed to adopt the 1996 Protocol to the London Convention, whose Annex 1 include "dredged materials" as matter that can be | https://www.epa.gov/sites/default/files/2015-10/documents/lpamended2006.pdf |

| | | | | |
|---|---|---|---|---|
| | | | "dumped", i.e., deliberately disposed of at sea and thereby subject aggregates mining to the convention. | |
| UN Convention on the Law of the Non-Navigational Uses of International Watercourses | Transboundary watercourses | 1997 | The Convention on the Law of the Non-Navigational Uses of International Watercourses was adopted by the UN General Assembly on 1997. In Art. 27 it highlights that "Watercourse States shall, individually and, where appropriate, jointly, take all appropriate measures to prevent or mitigate conditions related to an international watercourse that may be harmful to other watercourse States, whether resulting from natural causes or human conduct, such as flood or ice conditions, water-borne diseases, siltation, erosion, salt-water intrusion, drought or desertification". The removal of sediment through dredging and aggregates mining can lead to transboundary impacts, included in the provisions of this Convention. | https://legal.un.org/ilc/texts/instruments/english/conventions/8_3_1997.pdf |
| CBD First Strategic Plan on Biodiversity | Threats to biodiversity | 2002 | In decision VI/26 the Conference of the Parties to the CBD adopted a Strategic Plan for the Convention on Biological Diversity in 2002, in The Hague (The Netherlands). The COP urged Parties, States, intergovernmental organizations and other organizations to review their activities, especially their national biodiversity strategies and action plans in the light of the Strategic Plan for the CBD. | https://www.cbd.int/decision/cop/?id=7200 |
| SEA Protocol | Mining | 2003 | The Protocol on Strategic Environmental Assessment of the Espoo Convention (Convention on environmental impact assessment in a transboundary context adopted in 1991) was adopted in 2003. It establishes obligations of Parties to assess the environmental impact of major quarries, mining, and on-site extraction at an early stage of planning. The SEA Protocol augments the Espoo Convention by ensuring that individual Parties integrate environmental assessment into their plans and programmes at the earliest stages. | https://unece.org/text-protocol |
| ICCM "No-go" commitment | Mining | 2003 | In 2003, the International Council on Mining and Minerals (ICMM) made a "No-go" commitment for World Heritage sites. However, a growing number of World Heritage Sites are threatened by planned or active mining including aggregate and limestone mining. | https://www.icmm.com/en-gb/our-principles/position-statements/protected-areas<br>https://edepot.wur.nl/278977<br>https://whc.unesco.org/en/decisions/4478/<br>https://whc.unesco.org/en/extractive-industries/ |

| | | | | |
|---|---|---|---|---|
| Berlin Rules on Water Resources | Natural resources of rivers | 2004 | The Berlin Rules on Water Resources is a non-binding instrument adopted by the International Law Association to summarize international law customarily applied in modern times to freshwater resources, whether within a nation or crossing international boundaries. Art. 6 indicate that "States shall use their best efforts to integrate appropriately the management of waters with the management of other resources" and Art. 31 on the impact assessment process states that "Assessment of the impacts of any program, project, or activity shall include, among others: … c. Identification of ecosystems likely to affected, including an assessment of the living and non-living resources of the relevant water basin or basins". | http://www.cawater-info.net/library/eng/l/berlin_rules.pdf |
| Strategic Plan for Biodiversity 2011-2020 | Human pressures | 2010 | The 10th meeting of the Conference of the Parties, held on October 2010, in Nagoya, Aichi Prefecture, Japan, adopted a revised and updated Strategic Plan for Biodiversity, including the ***Aichi Biodiversity Targets***, for the 2011-2020 period (UNEP/CBD/COP/DEC/X/2). This is a ten-year framework for action by all countries and stakeholders to save biodiversity and enhance its benefits for people. | https://www.cbd.int/doc/decisions/cop-10/cop-10-dec-02-en.pdf |
| UN Declaration - The future we want | Mining | 2012 | The "Future We Want" is the declaration on sustainable development and a green economy adopted at the UN Conference on Sustainable Development, in Rio de Janeiro in June 2012, which highlights the role of ecosystem restoration in achieving sustainable development. Arts. 227 and 228 of the Declaration refer to mining: "… We further acknowledge that mining activities should maximize social and economic benefits, as well as effectively address negative environmental and social impacts. In this regard, we recognize that Governments need strong capacities to develop, manage and regulate their mining industries, in the interest of sustainable development"(Art. 227); "We recognize the importance of strong and effective legal and regulatory frameworks, policies and practices for the mining sector that deliver economic and social benefits and include effective safeguards that reduce social and environmental impacts, as well as conserve biodiversity and ecosystems, including during postmining closure. We call on governments and businesses to promote the continuous improvement of accountability and transparency, as well as the effectiveness of the relevant existing mechanisms to prevent the illicit financial flows from mining activities." (Art 228). | http://www.uncsd2012.org/content/documents/727The%252520Future%252520We%252520Want%25252019%252520June%2525201230pm.pdf%2520%2520 |
| Decision 37 COM 7 urging all WHC | Mining | 2013 | Decision 37 COM 7 on Emerging trends and general issues of the World Heritage Committee. In point 8 it highlights that "Notes with concern the growing impact of the extractive industries on World Heritage | https://whc.unesco.org/en/decisions/5018/ |

| | | | | |
|---|---|---|---|---|
| Parties to respect the ICMM "No-go" commitment | | | properties, and urges all States Parties to the Convention and leading industry stakeholders, to respect the "No-go" commitment by not permitting extractives activities within World Heritage properties, and by making every effort to ensure that extractives companies located in their territory cause no damage to World Heritage properties, in line with Article 6 of the Convention". | https://whc.unesco.org/archive/2013/whc13-37com-20-en.pdf |
| 2030 Agenda for Sustainable Development | Non-metallic minerals | 2015 | In 2015, the UN adopted the Sustainable Development Goals (SDGs) of the 2030 Agenda for Sustainable Development, which set out a 15-year plan to achieve 17 Goals. SDG 12 "Responsible consumption and production" is directly related to the supply of construction minerals as includes indicator 12.2.1 to register information on material footprint including the footprint of non-metallic minerals, whose main share corresponds to construction minerals. | https://documents-dds-ny.un.org/doc/UNDOC/GEN/N15/291/89/PDF/N1529189.pdf?OpenElement |
| WCC Resolution on "Protecting marine environments from mining waste" | Mining waste | 2016 | In 2016, the IUCN World Conservation Congress adopted the Resolution on "Protecting coastal and marine environments from mining waste" (WCC-2016-Res-053), which affirms that to meet Target 14.1 of the SDGs, and the objectives of UNCLOS and the London Convention and Protocol, regulations should be put in place to regulate and ultimately stop the use of marine disposal of mining waste (including dredged material). | https://portals.iucn.org/library/sites/library/files/resrecfiles/WCC_2016_RES_053_EN.pdf |
| WCC Resolution on "Avoiding extinction in limestone karst areas" | Limestone karst | 2016 | In 2016, the IUCN World Conservation Congress adopted the Resolution on "Avoiding extinction in limestone karst areas" (WCC-2016-Res-063) that urged governments, NGOs, academia, and industries to work collectively towards identifying and protecting hotspots of endemism and diversity in limestone karst areas and encouraged further research on the sustainable management of karst areas. | https://portals.iucn.org/library/sites/library/files/resrecfiles/WCC_2016_RES_063_EN.pdf |
| IPBES Assessment on Land Degradation and Restoration | Minerals including construction minerals | 2018 | See main text and Appendix S4. | |
| UNEA4 Resolution on Mineral Resource Governance | Minerals including sand | 2019 | In 2019, the 4th United Nations Environment Assembly (UNEA-4) adopted the Resolution on "Mineral Resource Governance" (UNEP/EA.4/Res.19), which recognizes the findings of the report "Sand and Sustainability: Finding New Solutions for Environmental Governance of Global Sand Resources" by UNEP/GRID-Geneva (2019) and encourages multiple actors to promote awareness of how the extractive industries can cause negative impacts on the environment if | https://wedocs.unep.org/bitstream/handle/20.500.11822/28501/English.pdf?sequence=3&isAllowed=y |

| | | | | |
|---|---|---|---|---|
| | | | not properly managed, and the sustainable mining and sourcing of raw materials to move towards decoupling economic growth from environmental degradation. | |
| UNE4 Resolution on Sustainable Infrastructure | Construction minerals | 2019 | In 2019, the UNEA-4 adopted the Resolution on "Sustainable Infrastructure" (UNEP/EA.4/Res.5). The UNEA invited Member States to further develop and implement sustainable urban development policies that promoted resource efficiency and resilience and respectively align sectoral policies such as transport, energy, waste management and sustainable buildings and construction, and encouraged all countries to promote public procurement practices that were sustainable, in accordance with national policies and priorities, stressing the importance of sustainable infrastructure for improving resource efficiency and sustainability in consumption and production processes and reducing resource degradation, pollution and waste, and fostering resilience to climate change, natural disasters and extreme weather events. | http://wedocs.unep.org/bitstream/handle/20.500.11822/28470/English.pdf?sequence=3&isAllowed=y |
| UN Decade on Ecosystem 2021-2030 | Human activities | 2019 | On 1 March 2019, the UN General Assembly adopted the resolution A/RES/73/284 and declared the UN Decade on Ecosystem Restoration. Through this, governments have recognized the need to prevent, halt and reverse the degradation of ecosystems worldwide for the benefit of both people and nature. The 2021–2030 timeline underlines the urgency of the task for also achieving the climate targets of the Paris Agreement or the Sustainable Development Goals. | https://wedocs.unep.org/bitstream/handle/20.500.11822/36251/ERPNC.pdf |
| IPBES Global Assessment Report | Minerals including construction minerals | 2019 | See main text and Appendix S4. | |
| WCC Resolution on "Reducing the impacts of the mining industry on biodiversity" | Mining | 2020 | In 2020, the IUCN World Conservation Congress adopted the Resolution on "Reducing the impacts of the mining industry on biodiversity" (WCC-2020-Res-121), which expresses concerns by the increased demand of minerals worldwide including construction minerals invites states and authorities to implement transition plans to reduce demand for primary materials and to phase down and progressively phase out their production and instead to supply recovered, reused and recycled materials and to find renewable substitutes, and calls on states to apply the precautionary approach to the management of risks to ecosystems from mining activities. | https://portals.iucn.org/library/sites/library/files/resrecfiles/WCC_2020_RES_121_EN.pdf |

| | | | | |
|---|---|---|---|---|
| WCC Recommendation on “For the urgent global management of marine and coastal sand resources” | Coastal aggregates | 2020 | In 2020, the IUCN World Conservation Congress adopted the recommendation “For the urgent global management of marine and coastal sand resources” (WCC-2020-Rec-029) motivated by serious environmental and social impacts of unsustainable sand mining in French Overseas Territories and recommending the implementation of strategic plans for the management of terrestrial and marine sand, taking appropriate measures for restoration, and systematically requesting impact assessments. | https://portals.iucn.org/library/sites/library/files/resrecfiles/WCC_2020_REC_029_EN.pdf |
| WCC Resolution on “Conservation of the natural diversity and heritage in mining environments” | Mining | 2020 | In 2020, the IUCN World Conservation Congress adopted the resolution on the “Conservation of the natural diversity and natural heritage in mining environments” (WCC-2020-Res-088), which calls Member States to conserve mining environments (open-cast mines and quarries), whose value derived from the conservation of their natural heritage is considered greater than the value of their restoration and encourages initiatives to ensure that the natural heritage of these mining environments is used for biodiversity conservation, and also to promote scientific, educational, cultural and/or tourist purposes. | https://portals.iucn.org/library/sites/library/files/resrecfiles/WCC_2020_RES_088_EN.pdf |
| UNEA5 resolution on Environmental aspects of minerals and metals management | Minerals including sand | 2022 | In 2022, the 5th United Nations Environment Assembly (UNEA-5) adopted a Resolution on “Environmental aspects of minerals and metals management” (UNEP/EA.5/Res.12) that stresses the need for enhanced action to support the environmental sustainability management of minerals and metals. It requests the Executive Director, through the Global Resource Information Database (GRID-Geneva), to “strengthen scientific, technical and policy knowledge with regard to sand, and to support global policies and action regarding the environmentally sound extraction and use thereof”. | https://wedocs.unep.org/bitstream/handle/20.500.11822/39927/ENVIRONMENTAL%20ASPECTS%20OF%20MINERALS%20AND%20METALS%20MANAGEMENT.%20English.pdf?sequence=1&isAllowed=y |
| Kunming-Montreal Global Biodiversity Framework | Minerals including construction minerals | 2022 | See main text and Appendix S2. | |
